# Improved modeling of RNA-binding protein motifs in an interpretable neural model of RNA splicing

**DOI:** 10.1101/2023.08.20.553608

**Authors:** Kavi Gupta, Chenxi Yang, Kayla McCue, Osbert Bastani, Phillip A Sharp, Christopher B Burge, Armando Solar-Lezama

**Affiliations:** Department of Electrical Engineering and Computer Science, Massachusetts Institute of Technology, Cambridge, MA 02139; Department of Computer Science, University of Texas at Austin, TX 78712; Department of Biology, Massachusetts Institute of Technology, Cambridge, MA 02139; Department of Computer and Information Science, University of Pennsylvania, Philadelphia, PA 19104; Koch Institute of Integrative Cancer Research, Massachusetts Institute of Technology, Cambridge, MA 02139

**Keywords:** alternative splicing, genome interpretation, machine learning, neural network, RNA processing, RNA-binding protein, transcriptome analysis

## Abstract

Sequence-specific RNA-binding proteins (RBPs) play central roles in splicing decisions, but their exact binding locations and activities are difficult to predict. Here, we describe a modular splicing architecture that leverages *in vitro*-derived RNA affinity models for 79 human RBPs and the annotated human genome to produce improved models of RBP binding and activity. Binding and activity are modeled by separate Motif and Aggregator components that can be mixed and matched, enforcing sparsity to improve interpretability. Standard affinity models yielded reasonable predictions, but substantial improvements resulted from using a new Adjusted Motif (AM) architecture. While maintaining accurate modeling of in vitro binding, training these AMs on the splicing task yielded improved predictions of binding sites *in vivo* and of splicing activity, using independent crosslinking and massively parallel splicing reporter assay data. The modular structure of our model enables improved generalizability to other species (insects, plants) and to exons of different evolutionary ages.

## Introduction

Genomic sequences encode the form and function of organisms, and their interpretation is an important scientific goal. The complex gene architectures present in metazoans makes this particularly challenging. Non-coding segments, or introns, frequently interrupt the coding sequence in eukaryotic genes (with 10 or more per gene on average in mammals), requiring intron excision and ligation of the flanking exons by the RNA splicing machinery in order to assemble the mature protein-coding mRNA^1^. Thus, finding the boundary elements or splice sites that are recognized in this process, the RNA splicing code, is important in interpreting the genome, and understanding how these sites are recognized by the spliceosome is a longstanding puzzle in molecular biology.

As the human genome was being sequenced and assembled, gene prediction was a priority, and incorporating detailed models of RNA splicing features proved useful^2^. Models more specifically focused on various aspects of splicing rather than gene finding have since been developed. The splice site motifs were the first aspects to be modeled, typically by giving scores for short sequences as potential 5’ splice sites (5’SS) or 3’ splice sites (3’SS) – the sites recognized at the beginning and ends of introns^3–5^. However, these motifs were found to be insufficient to predict splicing patterns, even in the simpler case where all introns are short ^6^, implying that other features must broadly contribute to intron recognition. Subsequent efforts have focused on splicing regulatory elements (SREs) – short RNA segments that typically function by recruiting splicing regulatory factors (SRFs), RBPs that promote or inhibit assembly of core machinery at nearby splice sites^1^ (Figure 1). Incorporating known exonic SREs into a simple splicing simulator was found to substantially improve the predictions of a simple splicing simulator^7^.

**Figure 1.**
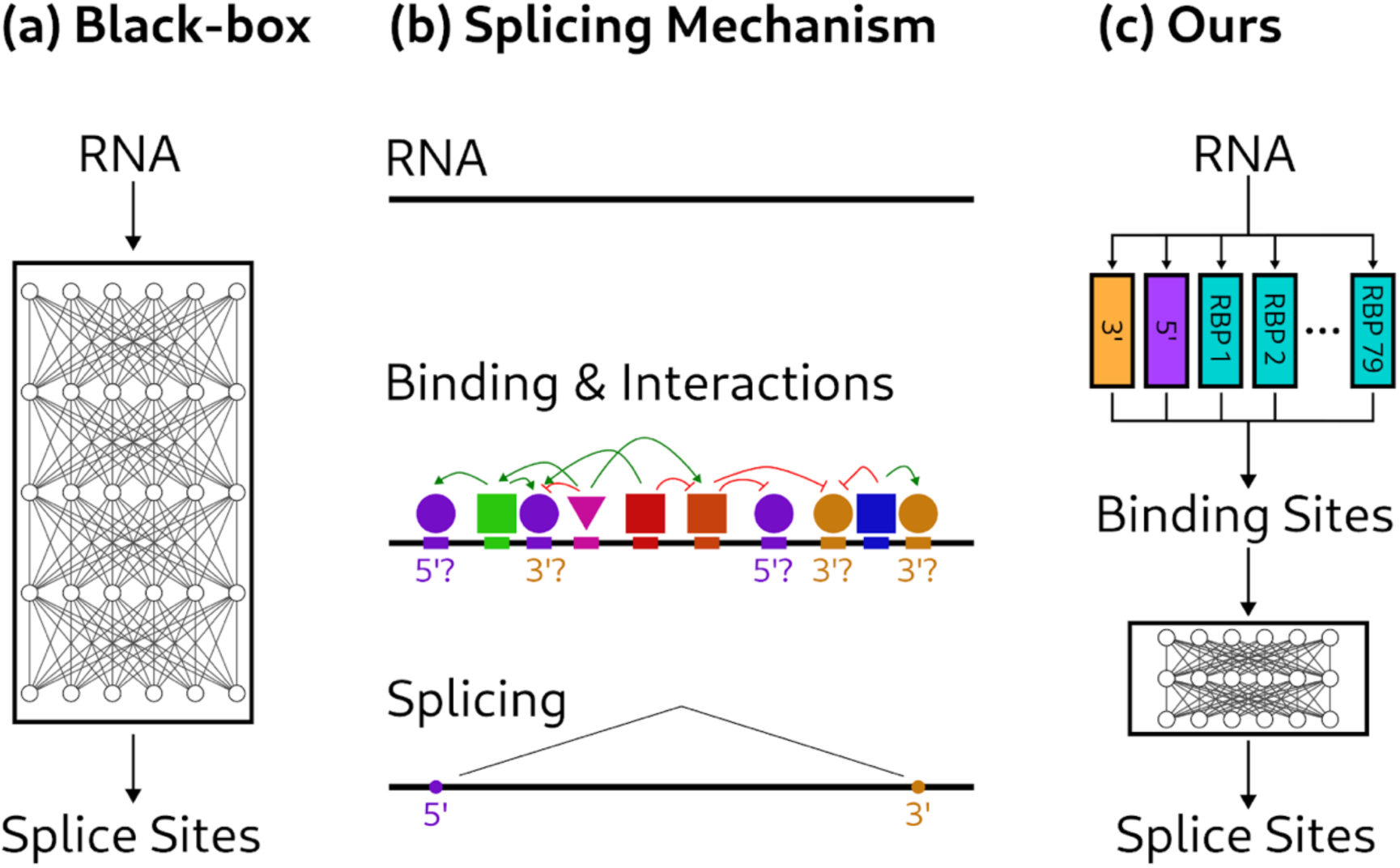
Types of models of RNA splicing. (a) SpliceAI models splicing as a black box mapping from pre-mRNA to splice sites. (b) We conceptualize splicing as a process in which various protein and ribonucleoprotein factors bind to the pre-mRNA sequence and influence which sites are chosen as splice sites and which are not. (c) Our SAM model predicts 5’ and 3’ splice site motifs and RBP binding motifs first, and then predicts which splice sites will be chosen by modeling interactions between these motifs.

Related problems have since been tackled, such as predicting the splicing phenotypes of mutations, or predicting aspects of alternative splicing such as the percent spliced in (PSI) values of exons, or the direction of changes in exon inclusion between different tissues^8–13^. Most models emphasized features known to be recognized in splicing, but to improve accuracy some models also considered features not available to the spliceosome, such as evolutionary conservation^14,15^.

More recently, some attention has returned to the question of predicting splicing from raw sequence using neural network methods that have been highly successful at predictive tasks in many fields^16–18^. Rather than curating features by hand, these machine learning methods have feature selection built into their training process, albeit in a manner that makes extracting and interpreting these features extremely difficult. For instance, SpliceAI, a deep convolutional neural network (CNN), is able to produce extremely high end-to-end accuracy in predicting splice sites in the human genome using up to 10,000 bases of sequence context. The model is essentially a black box that does not allow extraction of which specific features are of general utility in predicting splicing, although some clues can be gleaned about sequences important for prediction of individual splice sites using in silico mutagenesis^18^. Related CNNs have been developed for prediction of tissue-specific splicing patterns^19,20^. Some recent work^21^ has explored more interpretable learned models of splicing but has done so in the domain of short (70 nt) synthetic exons rather than gene-level splicing.

A recent line of research in AI has focused on interpretable neural networks that use some intermediate processing of the input that corresponds to known information about a particular process^22,23^. These techniques use auxiliary losses on intermediate layers in an effort to ensure that the network actually does make predictions based on general underlying data. Here, we instead use a hard constraint that requires the intermediate concept to be similar to previously known information, which is made possible using a sparsity-based approach. Other techniques use multiple datasets with the same intermediate features^24^, which is unfortunately not applicable to the splicing domain. Yet other techniques attempt to disentangle features of the input, in such a way as to extract information about the intermediate state^25^; however, these techniques attempt to preserve all information in the input, while we wish to force the model to use only a limited set of well-defined motifs and binding locations.

Here, we sought to develop splicing simulators that are interpretable and achieve high accuracy on gene-level splicing, with interpretability the paramount goal. This was achieved by developing sub-models inside of our Sparse Adjusted Motif (SAM) model that are directly anchored to known biology. Our Local Splice Site Identifier (LSSI) model learns the 3’SS and 5’SS motifs, and our Fixed Motif (FM) model estimates the binding locations of RBPs using biophysically-based models derived from in vitro RBP:RNA binding data^26,27^. We further developed an Adjusted Motif (AM) model which tunes these motifs as part of a general splicing prediction architecture. This approach affords a substantial increase in accuracy while ensuring that the AM models describe in vitro RBP binding comparably well as the original FM models, preserving the ability to infer the involvement of specific motifs and associated RBPs in splicing and to interpret variants that alter RBP binding.

## Results and Discussion

### A modular architecture for splicing with enforced sparsity

Our approach combines neural techniques with a modular structure that constrains the model to consider elements relevant to splicing. The three main model components are: 1) a Local Splice Site Identifier (LSSI) which models the 5’SS and polypyrimidine tract (PPT)/3’SS; 2) a “Motif Model”, which incorporates motifs representing in vitro binding specificity of several dozen RBPs; and 3) an “Aggregator”, which predicts the locations of splice sites based on the LSSI and Motif Model outputs (Figure 2).

**Figure 2.**
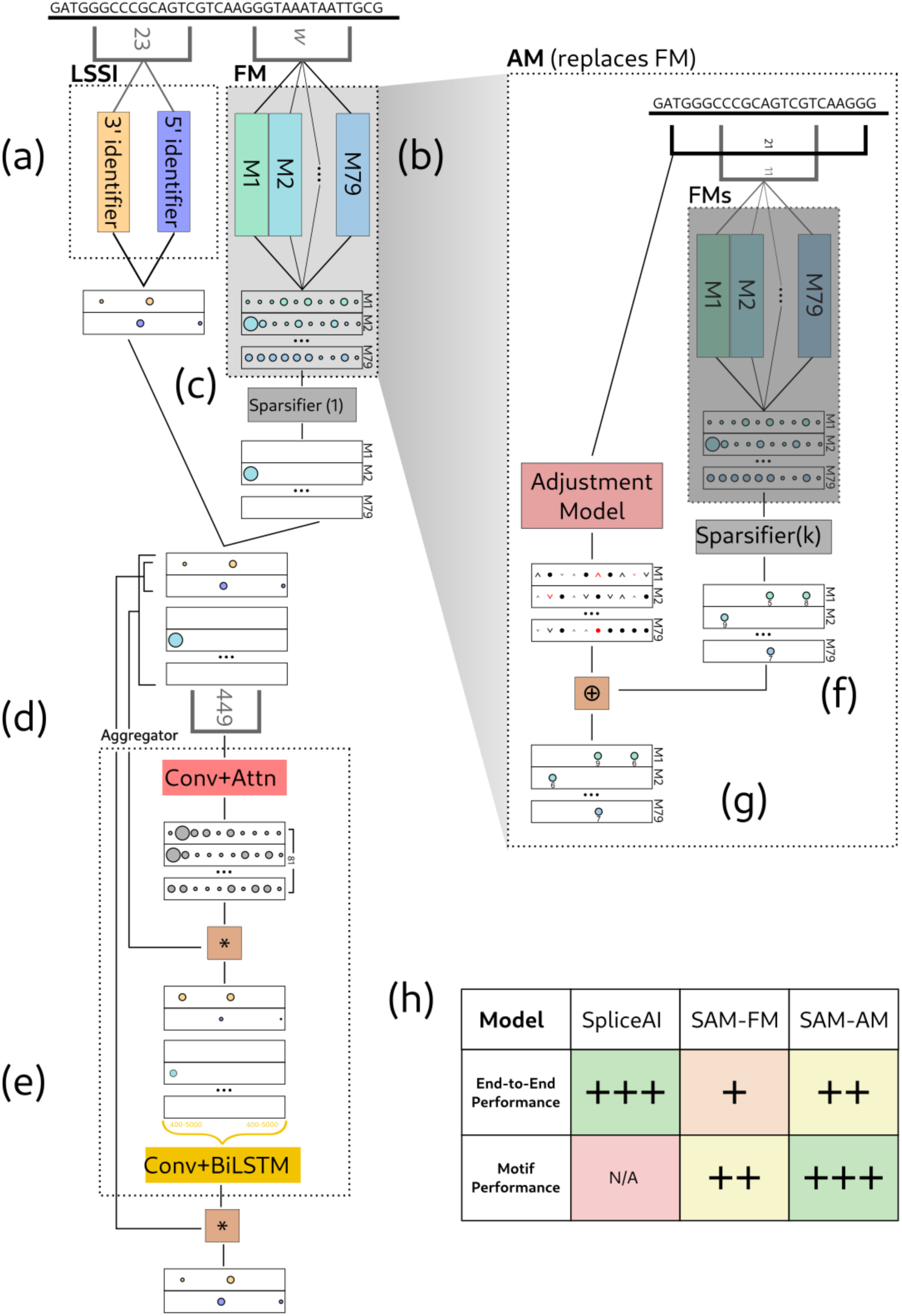
Overview of Sparse Adjusted Motif architecture: components and flow of information. Panels a) through e) summarize the “FM” model; panels (d) and (e) describe the “Aggregator” component of our model, inspired by ^31^; panels f) and g) summarize training of the “AM” models, which then replace the FM motifs shown in panel b). (a) The LSSI model processes the sequence and produces an annotation of the core 3’ and 5’ motifs. (b) The motif model processes the sequence and produces an estimate of RBP binding affinity at each site. (c) We enforce sparsity on the motif binding affinities, only allowing through high-affinity sites. (d) We compute influence scores for each position in the sequence; by multiplying with the sparse input we ensure that these influence values are only used to increase or decrease the strengths of known binding sites. (e) We then run a long-range processor across the sequence to score potential splice sites. We multiply these with the outputs of the core motifs to produce our final predictions. We use an LSTM here as the structure of an LSTM’s dataflow graph is identical to that of the forward-backward algorithm, which is commonly used to find the optimal sequence of states in an HSMM. (f) In our AM model, we first run our FM model and then sparsify it to a level of density *k* times denser than we intend to output (typically *k* = 2), and then derive AM parameters. (g) We also predict increase/decrease scores at each position. These scores are then added only to sites that were plausible binding sites. We then resparsify the output. This allows changing both the magnitudes of the sites arbitrarily as well as changing which sites are selected, while guaranteeing that all the produced AM sites are among the sites scored highly by the FM. (h) Table represents a high-level view of the two metrics of accuracy we consider. Both the FM and AM models are novel, though under the FM model the Aggregator is novel while the motifs are previously described. Our AM model improves over the FM model in all features, trading some accuracy versus SpliceAI in favor of being able to predict relevant RBP binding positions, and is the best at binding motif prediction. N/A indicates that SpliceAI is not capable of binding motif prediction.

The 3’SS and 5’SS have fairly strong core motifs, which are often captured by graphical models trained using maximum-entropy estimation or other methods^5^. We chose to model the same core splice site regions modeled by the MaxEnt method to facilitate comparison. The 3’SS region comprises 23 nt, consisting of the last 20 bases of the intron and first 3 bases of the succeeding exon, while the 5’SS region comprises 9 nt, consisting of the last 3 bases of the exon and first 6 bases of the downstream intron. In general, our LSSI models slightly outperform MaxEnt models, with top-*k* accuracies for 3’SS and 5’SS on the SpliceAI test set of 23.5% and 26.5% versus 18.4% and 24.7% for the MaxEnt model. (Top-k accuracy is the fraction of splice sites predicted correctly at a score cutoff which predicts the same number of splice sites as actually appear in the data.)

Next, we considered models of RBP binding sites in the Motif Model. In our baseline model, we use what we call Fixed Motifs (FMs), which consist of position-specific affinity matrices (PSAMs) derived by the RBPAmp algorithm^27^ from RNA Bind-n-Seq (RBNS) in vitro data for 79 human RBPs^26^. Each RBP is represented by 1-5 PSAMs (typically ~11 nt long) that yield relative affinity values for any RNA sequence. We have also explored using PSAM models inferred from RNACompete data^28^. To score a sequence with the PSAM(s) for an RBP, we compute relative affinity from the matrix, then multiply this value by the corresponding absolute affinity, and sum the affinities across all PSAMs. Our primary contribution, the AM Model described below, is an alternative to the PSAM-based FM model.

One consideration in designing a modular neural network structure is to ensure that the LSSI and Motif model outputs aren’t simply used by the Aggregator to reconstruct the original sequence, and then learn a black box splicing algorithm. If this were possible, high accuracy might be achieved without the essential properties of modularity and interpretability. Therefore, it is important to ensure that these models compress the information in the sequence, thus preventing it from being reconstructed. Enforcing an information bottleneck in order to ensure modularity is a standard technique in machine learning^29^; we use a recently-described approach^30^.

For the LSSI model, we ensure sequence compression by enforcing the minimum score to be a log-probability of –10, by setting all values below –10 to –10. This bar, if interpreted as a binary classification bar, has the effect of including almost all splice sites (98.88% of 3’SS, 99.10% of 5’SS) while assigning meaningful (> –10) scores to just a small fraction of locations (2.00% for the 3’SS model, 1.41% for the 5’SS model, a mean of 1.7% of positions). The LSSI model emphasizes recall over precision as one of its functions in the model is as a mask: any position scored –10 by both LSSI models will not be considered as a potential SS subsequently. Since we assign meaningful values to only 1-2% of locations for each splice site type, and this model is trained entirely separately from the main model, it greatly compresses the input sequence and could contribute at most minimally to sequence reconstruction.

### Enforcing Motif Sparsity

When modeling SRFs, it is more important to precisely bound the mutual information between the input RNA sequence and the motif layer, as there are many RBPs and thus much more information could theoretically pass through. We use the fact that we know that RBPs bind specific sequences and have finite free concentrations, so that only a fraction of sites in the transcriptome are occupied by RBPs. Therefore, we add a layer immediately after our Motif Model that enforces sparsity, limiting the fraction of sites at which the model considers binding by an RBP. We enforce the density, the number of positions that contain a nonzero value, to be at most δ by enforcing that the sparsity is at least (1 – δ). If we have *M* motif models, if each predicted RBP binding score contains at most *η* bits of information, and we analyze *L* bases, we can bound the entropy of the motifs layer by *H/L* ≤ *M[H(B(δ)) + δ η]*, where *B(x)* represents the Bernoulli distribution with probability *x*, *δ* is the mean motif density, *and η* represents an upper bound across channels of the entropy in the distribution of nonzero activations (Supplementary Figure 1; derivation provided in Supplementary Text).

This quantity represents the information per base that is being let through by the Motif Model. In general, we require *H/L* < 1.91 bits, in order to ensure that the sequence is being compressed. Throughout this paper, we use *M* = 79, which is the number of RBPs for which we used PSAMs from RBNS as input, unless otherwise specified, and we set *δ* = 0.18% to be our definition of sequence compression. This value allows *η* < 2.8 bits while maintaining sequence compression. In general, the outputs of our sparse layer contain less than 2.8 bits each, with typical values of about 2 bits (Supplementary Text).

We empirically confirmed that our entropy bound is sound, but not particularly tight, via an experiment where we trained a neural network to reconstruct the sequence. From the motif output of an AM model with *H/L* bound of 1.79b/nt the original transcript sequence could be reconstructed with only 59% accuracy in one experiment. This corresponds to an effective *H/L* of 1.16 bits/nt. However, we continue to use 0.18% density in order to ensure that the motif density is well below the theoretical limit required to guarantee compression. Enforcing an entropy bound therefore ensures that the algorithm learns a compressed representation of the sequence and cannot reconstruct the input sequence.

To enforce a maximum density of *δ*, we follow the Sparling technique^30^, with a minor variation that we find improves accuracy at the cost of training efficiency. To train this model, our “standard training” approach is as follows. We start with a very large *δ* value of 75%. As training progresses, we reduce the maximum density threshold *δ* by a factor of 0.75 at particular steps, allowing the thresholds to update via the moving average. Rather than pick a certain number of training steps between reductions in *δ*, we instead use validation accuracy as a guide. At specific steps *t* (20 times per epoch), we check whether *V(t)* has reached some target accuracy *VT(t)* and, if so, reduce *δ* to 75% of its previous value.

Unless otherwise specified, we choose *VT(t) = VT* to be a constant in *t*, and tune the value of *VT* based on a search for *VT* such that we can train our model to achieve a density of *δ*=0.18%. In some experiments where it was of interest to evaluate the accuracy resulting from various minor model or data changes, we instead use a “quick training” method equivalent to the Sparling approach. In this approach, we set *VT(t)* to a dynamic function that starts at a high value (typically 85%), then reduces by 1% per epoch of training, increasing to whatever validation accuracy was achieved at the previous step whenever δ is reduced. Though faster, the quick training approach tends to produce slightly (~1-2%) lower performance.

We use the same train/test split as SpliceAI, with the following modification: we treat the first 50% of the SpliceAI test set (by total sequence length) as a validation set, which is used to determine when to adjust sparsity/accuracy thresholds. We use the last 45% of the SpliceAI test set as a true test set, for our evaluation results, with the 5% gap serving to avoid overlap of genes between training and validation sets. Different random orders of genes are used in training/testing.

### Aggregator

The final component of SAM is the Aggregator, a network that aggregates potential splice sites and RBP binding sites identified by the LSSI and Motif Model and produces a prediction of splice site locations. This network is structured in the following way: the LSSI output is concatenated with the Motif model output, and then processed by a small convolutional model of width 49 nt, then an attention layer of width 401 nt, which is then multiplied with the output of the motif model to propagate the sparsity. We use a custom attention layer design that takes advantage of this limited attention region for performance reasons. We then apply another CNN followed by a long-range BiLSTM network, and the result is then multiplied by the output of the LSSI model to produce our final splicing prediction. The design of the Aggregator is inspired by the hierarchical attention network structure^31^. However, without the multiplication and convolution (pre-BiLSTM) operations, the model cannot effectively capture the intrinsic importance of the motifs. We deliberately decided not to focus on or optimize the Aggregator in this paper, mostly treating it as an opaque box, except in a few instances where we use it to infer patterns of motif activity. We have not extensively investigated whether this exact architecture is optimal for our purposes, but we provide an analysis comparing our Aggregator with SpliceAI (Supplementary Figure 2).

### Adjusted Motif Model

We observed that the accuracy achieved with FM models was moderately high (67.2% of splice sites correct, see below for details) but wondered whether these models of in vitro binding of 79 RBPs were optimal for predicting splicing. As an experiment, we allowed the parameters to change subtly during training, while taking steps to preserve the association with individual RBPs, yielding “adjusted motifs” (AMs) (Figure 3). For example, binding preferences of RBPs in vivo might occasionally differ from the in vitro-derived FM models as a result of binding partners^32^ or post-translational modifications^33^. Furthermore, many RBPs belong to families of related proteins, usually with similar but not identical RNA-binding preferences, so the AMs might learn the binding preferences of the SRFs with most important roles in splicing from information about related family members analyzed in vitro. Training an unconstrained neural model to replace the FMs might also improve accuracy, of course, but at the expense of interpretability, as the learned motifs would no longer correspond to known RBPs.

**Figure 3.**
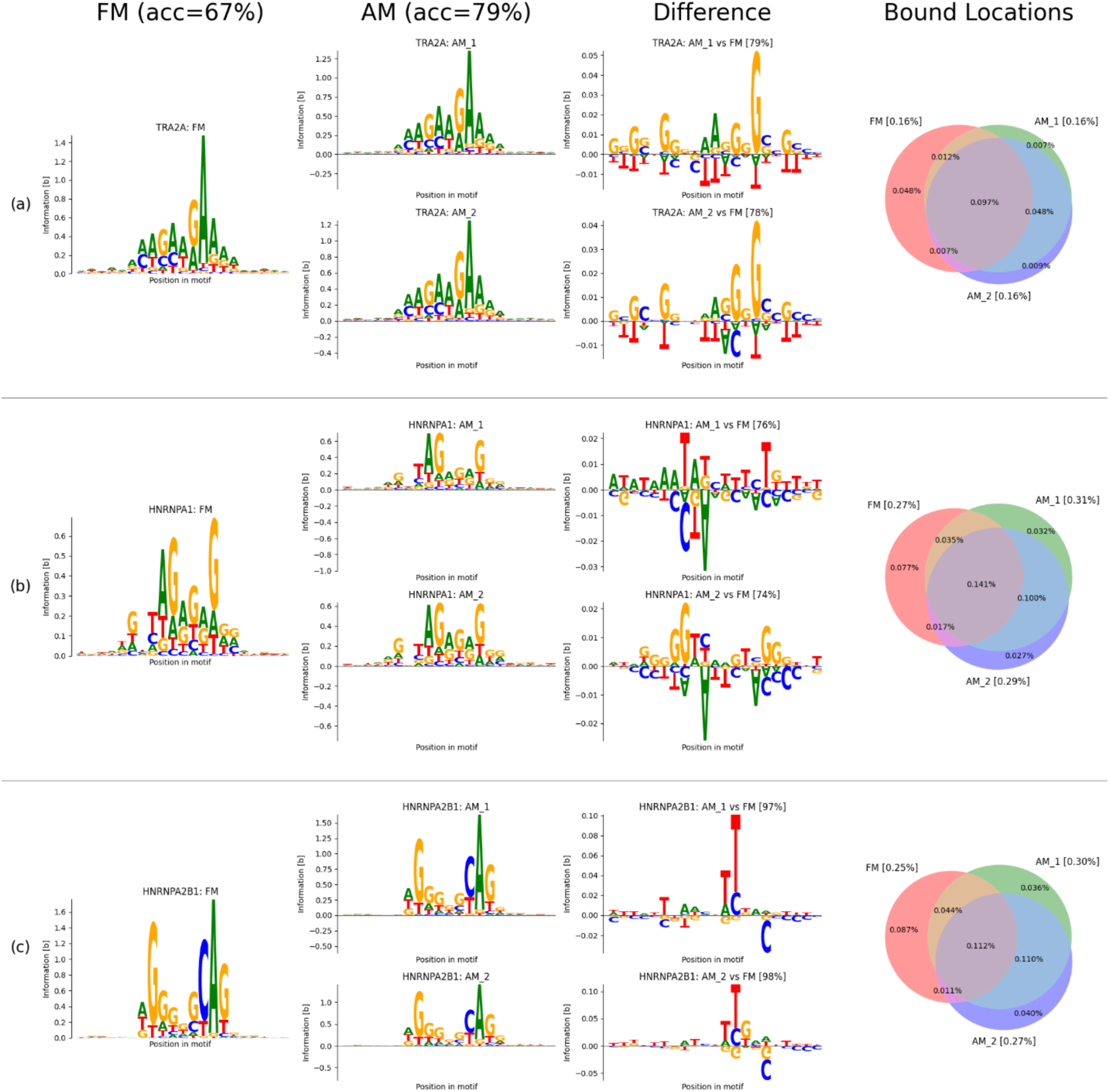
Sequence logos for corresponding AMs and FMs show overall similarity with minor differences. Sequence logos are shown for three representative SRFs: TRA2A, a splicing activator, and splicing repressors HNRNPA1 and HNRNPA2B1. The first column shows the FM motif and the second shows AM logos for 2 replicate runs. The third column shows differences between each AM and the corresponding FM: each base’s height represents its pointwise mutual information, with those above the bar enriched in the AM, and those below enriched in the FM. The pattern of differences is often but not always related between replicates, as shown by the examples above. The accuracy a logistic model can achieve at distinguishing AM from FM motifs is shown above each difference plot. The fourth column shows the overlap between FM and AM binding sites for each factor, labeled with the fraction of genomic positions for each category. The total fraction of genomic positions for each model is shown next to the model’s name in square brackets. For example, FMs and AMs for TRA2A bind 0.16% of genomic sites, i.e. one site every 625 nt.

We compute the binding sites in the adjusted motif (AM) model by adjusting the scores for already relatively high-scoring FM binding sites, allowing *k* times as many FM sites as we will eventually use, allowing our AM model to filter out a subset of binding sites among plausible sites provided by the FM model (Methods). Here, we have used *k*=2: for example, the top 1000 sites scored by an AM model would be constrained to fall within the top 2000 sites scored by the corresponding FM model. This value of *k* serves to constrain the sequence features of AMs to closely resemble the FM motifs in virtually every case, as desired (examples are shown in Figure 3). Each AM was modeled as a CNN composed of 5 residual units, each with two convolutional layers of width 3, resulting in a total context window of 21 nt. An exploration of motif widths obtained better results with width 21 than with shorter widths (Supplementary Figure 3).

We also directly examined the sequence logos corresponding to each motif to assess the adjustments. In general, we found that the motifs tend to be quite similar, with slight differences at a subset of positions (Figure 3), and modest variation from replicate to replicate. To assess the information captured by the AM models, we trained a logistic classifier with four parameters corresponding to every position in the motif (one-hot encoding of the nucleic acids) to predict the difference between sites classified as bound exclusively by the AM and those classified as bound just by the FM. In general, these classifiers had 75-100% accuracy (average of 90% across RBPs), demonstrating that a large fraction of the differential information captured by the AMs is attributable to linear combinations of the bases present at different motif positions.

The substantial improvement in splicing prediction from the AM models, while maintaining discrimination in the in vitro binding task, raised the question of whether neural models specifically trained on the RBNS dataset could improve splicing prediction (See section Methods: Neural models trained on RBNS data for more references). The FM models have widths of ~11 nt, but a neural model with width 11 nt did not improve discrimination on the RBNS dataset (–0.02% average change in performance), suggesting that the PSAMs are near-optimal representations at width 11. A neural model of width 14 nt, and an AM-architecture model (of width 21) trained on RBNS data improved performance on RBNS data by 0.33% and 1.76%, respectively. We refer to the 21-wide AM-architecture model trained on RBNS as a “Neural Fixed Motif” (NFM) model, since the parameters are frozen after training on the in vitro RBNS data and not allowed to vary during training on the splicing task. When the NFM was used in splicing prediction with an adaptive accuracy threshold, performance was 64.1%, slightly below the FM model’s 65.3%, and far below the accuracy of 78.6% achieved by Ams trained on the splicing task (these values are lower than those presented earlier as they resulted from the quick training approach). Our interpretation of these observations is that training of AMs on the splicing task using genomic data enables learning of specific sequence features relevant to binding and/or splicing in vivo that are likely not present in the in vitro RBNS data. For example, the AMs might be learning a motif bound in vivo that differs somewhat from the in vitro motif because of the presence of specific post-translational modifications or binding partners of the RBP. Alternatively, instead of the motif of the single RBP analyzed in vitro, the AMs might be learning a motif bound by a close paralog, or a composite motif bound by some or all members of the protein family to which the RBP belongs, and/or other proteins.

As a control, we checked to see to what extent RBNS binding activity correlated with positions that SpliceAI found important. We performed an in-silico mutagenesis experiment on SpliceAI, computing an activity score for positions in the sequence as the effect that mutating them had on SpliceAI score. While 3’SS and 5’SS sites clearly had more of an effect than other sites, this was not true of RBP binding sites as predicted by RBNS. On the other hand, we found that for SAM models, both FM and AM, RBP binding sites did tend to have greater activity scores (See Supplementary Figure 4 for more information). We can thus conclude that SpliceAI is not representing a model of splicing equivalent to ours, and is focusing instead on other features, which may or may not correspond to RBPs.

### Module Substitution Experiment

Our SAM model is designed to be explicitly modular. Setting aside the LSSI for now, which was unchanged during training, our model can effectively be considered as a composition of a Motif Model, *M,* and an Aggregator, *A*. In a traditional neural network, the intermediate layers of a model represent latent variables that are not necessarily stable across different models, as they may converge to different representations of the same information or may even represent different pieces of information relevant to the learned task. However, in our case, we intend M and A to represent the non-latent concepts of RBP binding sites and the activities of these RBPs in splicing, respectively. If this expectation holds, we should be able to use one model’s *M* with another model’s *A* and still achieve good performance. An alternative, undesirable possibility would be if the AM models side-channel information through their outputs that the Aggregator then learns to pick up, but which do not represent activities of the RBPs.

To distinguish between these alternatives, we conducted a computational “Module Substitution” experiment (Figure 4) in which we use the AM models as the motif model in our architecture and combine it with the Aggregator trained to work with the FMs. One subtlety here is that our training process for AMs cannot guarantee that the actual values of activations are similar across models, so we resolve this by adding a “binarization” layer to pre-trained models that sets all non-zero motif binding site values to 1, followed by retraining of the Aggregator.

**Figure 4.**
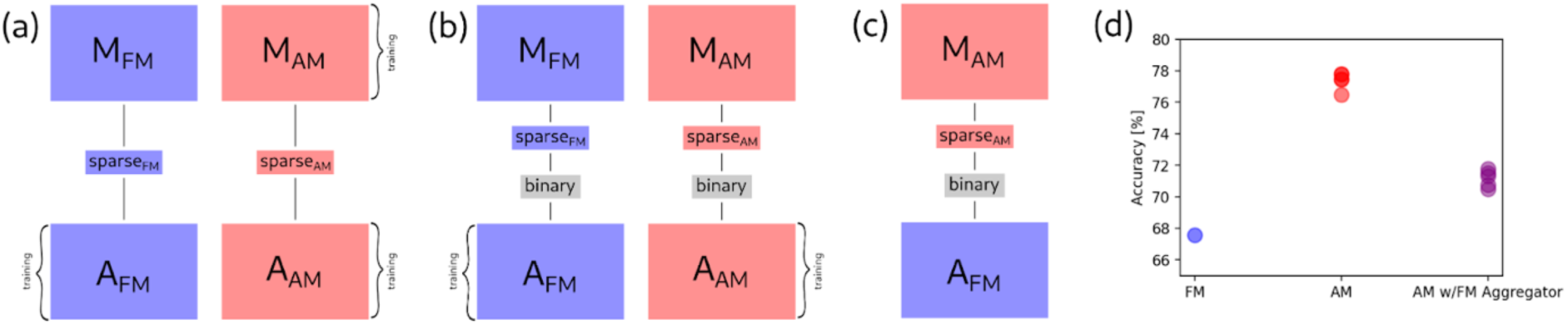
Module substitution shows that Aggregator learns similar SRF activities when trained with FMs or AMs. In this experiment, we demonstrate that the AM model still produces motif sites that carry the same semantic meaning as the original FM sites. To this end, we take the AM motifs and send them to the FM aggregator. If the AM model were producing non-motif information, this would lead to a degradation or at best no change in performance. However, we observed an improvement, from ~68% to ~71% accuracy, demonstrating that the AM models are better at predicting motif binding sites, even provided to a model trained to work with FM predictions. **(a-c)** Stages of the module substitution process. The FM Motif Model and Aggregator are in blue, while those corresponding to the AM model are in red. (a) Both models are trained as usual. (b) The models are binarized by applying a binary layer and then the FM model’s Aggregator is retrained to handle the new binary motif outputs. (c) The new FM Aggregator is used in conjunction with the binarized AM motif model, with sparse layers and binarization layers as shown. (d) Results of the combination experiment.

The results of this experiment (Figure 4) reveal two clear trends. Comparing to our previous results (Figure 3), the AM models tend to perform slightly (1.2-2.9%) worse after binarization, while the FM models are unaffected (Supplementary Figure 1). This observation suggests that the AM models provide information through motif “strength” as well as location, while the FM models’ utility is primarily in identifying potential locations of binding. More importantly, AMs tend to outperform FMs across the board, even when using the FM Aggregator. Since the FM Aggregator was trained without ever seeing AMs, it cannot be side-channeling information unique to the AMs. Instead, the consistent improvement seen with AMs here demonstrates that they largely provide information of the same type (i.e. binding of the same RBPs) as provided by FMs, but of improved quality. The further improvement seen when using the AM Aggregator is expected and may reflect that this model makes better use of the motif strength information provided by AMs.

### Analysis of eCLIP binding data

To understand how well the FM and AM models reflect binding conditions in vivo, we used data from the enhanced crosslinking and immunoprecipitation (eCLIP) assay, which has been widely applied to human RBPs in two cell types as part of the ENCODE project^34^. The eCLIP datasets consist of sets of eCLIP peaks, each consisting of a start and end location in the genome that represent the approximate binding location of the associated RBP. At no point were our AM or FM models trained on eCLIP data in any way, so these data provide an independent assessment of the extent to which these models capture RBP binding information. All our analysis results in this section were performed on the subset of 18 RBPs where we had both RBNS PSAMs and eCLIP peak data. We defined “eCLIP regions” by extending the eCLIP peak an additional 50 nt to the 5’ end, since this was found to enrich for known binding motifs^35^. We selected a random sample of 10,000 5 kb regions from the SpliceAI test set for our experiments.

Our metric for eCLIP accuracy is eCLIP enrichment of a model versus the FM model at equivalent sparsity, calculated separately on intronic and exonic sites. It was important to control for intron/exon bias as eCLIP tends to preferentially detect exons, both because some crosslinking may occur in the cytoplasm which is essentially devoid of introns, and because even in the nucleus introns have shorter half-lives than exons because they are rapidly degraded after excision. Thus, any bias toward exons could skew the results. Therefore, we present results independently for introns and exons. It was also important to use equivalent sparsity, since enrichment is sensitive to small changes in sparsity (Methods).

As a point of comparison, we also trained AM-format models directly on eCLIP data, designating these models “AM-E”. We did so by setting up a training objective where the model’s output was treated as predicting the probability that each location within a peak is not a match (treating the AM output as the negative log probability of non-match), then multiplying those probabilities together to produce an overall probability of non-match in each peak, then using the logistic loss against whether or not the match is a control peak. We balanced the dataset to be 50% intron and 50% exon, sourcing sequences from the SpliceAI training dataset. As they were specifically trained to solve the eCLIP enrichment task, these AM-E models provide an upper bound on the performance one might expect any model to achieve on this task, though this bound is not necessarily tight as eCLIP-trained models may also learn technical eCLIP biases (e.g., related to the efficiency of crosslinking of different sequences), as well as true RBP binding signal. When used as part of an end-to-end model of splicing, the AM-E models underperformed FM models using the same 18 motifs, achieving an accuracy of 50.3% vs 56.8% for the FMs, suggesting that FM models better represent RBP binding sites relevant to splicing.

The results of these experiments (Figure 5a) show that, across several seeds, our AM models improve on FMs in prediction of eCLIP binding, achieving about 50% of the theoretical possible improvement on exons, and 10% on introns. These observations indicate that, in the process of modeling splicing end-to-end on the genome, the AM models have learned to better model the binding locations of RBPs in vivo. This observation supports that the constraints built into our model have encouraged modeling of biochemical features relevant to splicing.

**Figure 5.**
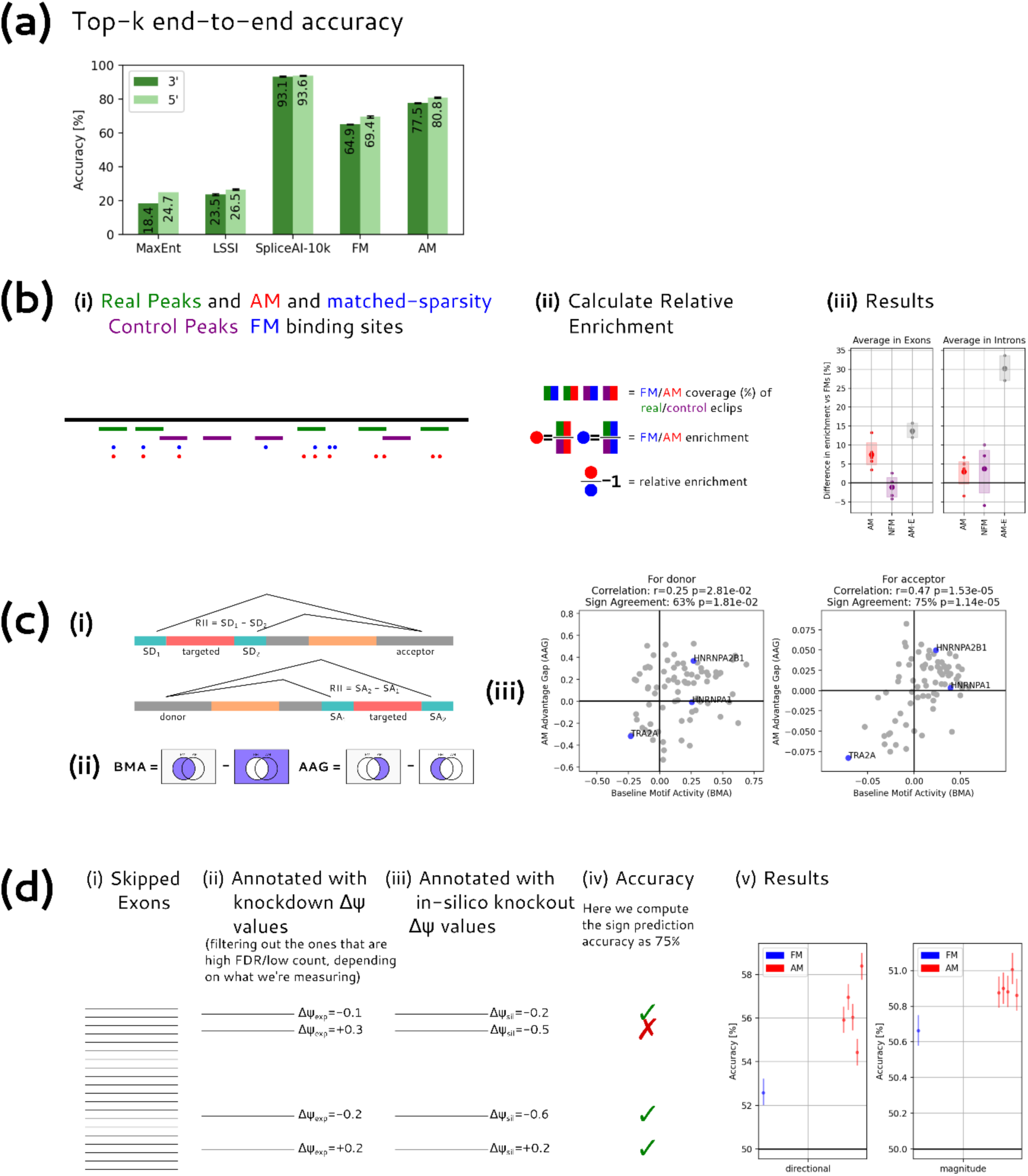
AM motifs perform better at predicting splicing, eCLIP peaks, MPRA activity, and knockdown data than FM motifs. The SAM model performs improves substantially at end-to-end prediction over existing RBNS-derived FM motifs, though it underperforms SpliceAI. In addition, the AM model, unlike SpliceAI, can also perform other motif-specific tasks, in all cases better than the FM motifs. **a) End-to-End Accuracy:** Bars represent top-k accuracy of prediction of 3’SS (dark green) and 5’SS (light green) for different models trained on the SpliceAI training set and tested on the last 45% of the SpliceAI test set (the rest is used for validation. This 45% constitutes 37.14M nt across 730 genes). Error bars indicate the minimum and maximum performance among 5 replicates. The MaxEnt model is deterministic so does not have an error bar. **b) eCLIP peak prediction** (i) Schematic of a genomic region showing real eCLIP peaks (green) and randomly generated control peaks (purple). The control peaks are generated by shifting the locations of peaks for a given RBP to random positions in the same transcript. AM motifs (red) and sparsity-matched FM motifs on the sequence are also shown. (ii) We count peak overlap for each of the four comparisons depicted (in practice, we collect this separately per motif and average across all 4. We compute enrichments for each model, and then calculate the % increase from one to the other. (iii) Plotted are the mean relative enrichments across motifs, separately for exon and intron. AM-E models are provided as a reference for the maximum improvement we could hope to achieve using a motif model on eCLIP data. By this standard, the AM models achieve about 10% of the theoretical maximum performance improvement over FMs on introns, and about 50% of the theoretical maximum performance improvement on exons. **c) MPRA activity prediction** (i) Schematic of the experimental setup from^36^. Reporters were used involving a pair of alternative 5’ (donor) splice sites, or a pair of alternative 3’ (acceptor) splice sites, such that the degenerate region in red is either included in the exon or spliced out (triangular lines depict possible introns). We ignore the other degenerate region (in orange). To compute the Relative Intron Inclusion (RII) score, we subtract the read count of the splice site that indicates the red region is in the intron from the one that indicates it is not. (ii) We compute the Baseline Motif Activity by looking at just FM sites and non-sites, as we are confident that FMs indicate binding. We compute the advantage gap for AMs by looking at the difference of uniquely AM and uniquely FM slices: note that this is a symmetric measurement. (iii) The relationship between these two metrics is significantly positive, on both the 3’ and 5’ data. We highlight the three well-established motifs from Figure 6 in blue. **d) Knockdown modeling** (i) we use an existing dataset of skipped exons and corresponding in vivo knockdown results, (ii) filtered for reliability both by looking at low FDR and by looking at high counts (iii) we run our own model to compute in-silico knockdowns. (iv) our metric of interest is predictive accuracy of using the in-silico values to predict the knockdown values, both using sign and magnitude above/below median (iv) results at predicting experimental sign and magnitude from in silico sign and magnitude. Different AM models indicate different replicates, error bars indicate 95% bootstrap confidence intervals.

### Splicing Activity Analysis

Next, we considered whether the AM models have captured information relevant to splicing regulation in vivo. For this purpose, we turned to data from a recent splicing massively parallel reporter assay (MPRA) experiment^36^, using the randomized segments between alternative 5’SS motifs and between alternative 3’SS analyzed in this study (Figure 5b). Since these regions are located in introns only when the distal splice point is used (SD1 or SA2), we can use the differences in splicing SD1 – SD2 and SA2 – SA1 as measures of much a sequence favors recognition as intron versus exon, referred to as the relative intronic inclusion (RII) of a given segment.

We consider a randomized segment to be bound by a certain RBP according to a motif model if the motif model predicts at least one binding site in the region. We did not use the Aggregator model in this experiment, since the sequences involved are far shorter than those handled by our model. Instead, we compute a baseline motif activity (BMA) for each motif by calculating the mean RII in segments that are RBP-bound according to the FM model minus the mean RII in segments not predicted to be bound. The BMA establishes an activity for an RBP as promoting recognition of bound regions as intron (if BMA > 0) or exon (if BMA < 0). To emphasize differences between the models, we compared sites bound only by the AM Model (AM \ FM), and sites bound only by the FM model (FM \ AM), with the difference in RII change referred to as the AM Advantage Gap (AAG). Assuming that the BMA reflects the true activity of a factor, if ASA and BMA have the same sign then the AM model better captures binding, while if they have the opposite sign then the FM better captures binding.

We calculated AAG versus BMA in 5’SS and 3’SS datasets, averaged across five runs (Figure 5b). For both categories, the correlation of the values and signs is positive, and statistically significantly (p < 0.05 for 5’SS, p < 0.0001 for 3’SS). (The presence of some non-SRFs in the 79-RBP set may add some noise to the results.) Thus, we can conclude that the AM models, in general, yield better models of splicing-relevant motif activity than FM models. In a similar experiment using NFMs, BMA and AAG had the same sign 46% and 41% of the time on the 5’SS and 3’SS datasets, respectively, i.e. no advantage over the FMs. Therefore, these observations support that it is the training of the AMs in the end-to-end splicing training task that has improved their prediction of binding associated with splicing regulation.

### Analysis of RBP Knockdown Data

To further explore the ability of our models to infer the effects of RBPs on splicing, we used the RNA maps dataset from the ENCODE project^34^. This dataset consists of skipped exons annotated with change in percent spliced in (“delta ψ”) values with RNAi knockdown of a specific factor in human K562 and/or HepG2 cells; here, data from the two cell lines were combined to increase statistical power. Specifically, as described in Figure 5, we compute “in silico” delta ψ values by first predicting ψ, treating the model’s probabilistic prediction of splicing as a ψ value, and then subtracting from this value the ψ resulting from an “in-silico knockout” where all occurrences of the relevant motif are zero-ed out (i.e. treated as absent) before passing the values to the Aggregator.

We use two separate targets in our experiment. First, we compute the proportion of exons for which the correct sign of delta ψ is predicted (Figure 5c), focusing here on exons with FDR < 0.05, i.e. those which showed a significant difference in ψ following knockdown. Secondly, we compute the accuracy of predicting the magnitude of experimental delta ψ from the magnitude of in-silico delta ψ, specifically predicting whether the magnitude is above or below the median of exons (filtering for experiments having at least 50 samples in both control and knockdown conditions).

This experiment has several limitations. First, our model has been trained entirely on “canonical” splice sites, which are mostly constitutive, therefore may not capture unique features of alternative splicing. Second, our in-silico knockouts are not a perfect match for the experimental system, in which the target protein level is depleted but not eliminated, and our model does not consider secondary (often compensatory) effects in which the expression of other SRFs changes in response to the knockdown. These considerations likely reduce the agreement between the in-silico and experimental knockdowns. Despite this, we find that in general, our models perform better than chance at predicting the sign and magnitude of delta ψ values, and that AM models outperform FM models. These observations support that our model captures aspects of the splicing regulatory activity of many RBPs, and that this regulatory activity is better modeled with AMs than FMs.

### Interpretability and Modularity

To better understand the inner workings of our model, we conducted a series of computational experiments aimed at understanding the regulatory logic learned by SAM models. We first used in silico motif deletion to assess the regulatory impact of different motifs on each model’s predictions of individual genomic regions, comparing the FM and AM models with LSSI as a baseline reference; a sample exon and sample decoy exon are shown in Figures 6a and 6b, respectively. For exon 54 of the *DNAH14* gene, the splice sites are too weak to be predicted by the LSSI model. However, motif deletions showed that the AM model predicts both splice sites correctly primarily because of positive impacts from RBM41 and TRA2A motifs located inside the exon. The FM model identifies some of the same motifs (e.g., the RBM41 motif), but not the TRA2A motif and considers additional negative-regulatory motifs, causing it to miss the exon. In the second example, a “decoy” (false) exon is predicted by the FM model in the gene *SNAP47*, with the 5’SS also predicted by the LSSI model, but the AM model correctly does not predict the decoy exon because of inhibitory effects of three RBP motifs (associated with SRSF8, FUS and HNRNPA0) not identified by the FM model. These examples were curated for compactness and to illustrate cases where AM and FM models differ, but such a ‘putative regulatory landscape’ can be generated for any exon or genomic region of interest, with uncertainty estimates possible using several different model seeds. Additional examples are shown in Supplementary Material.

**Figure 6.**
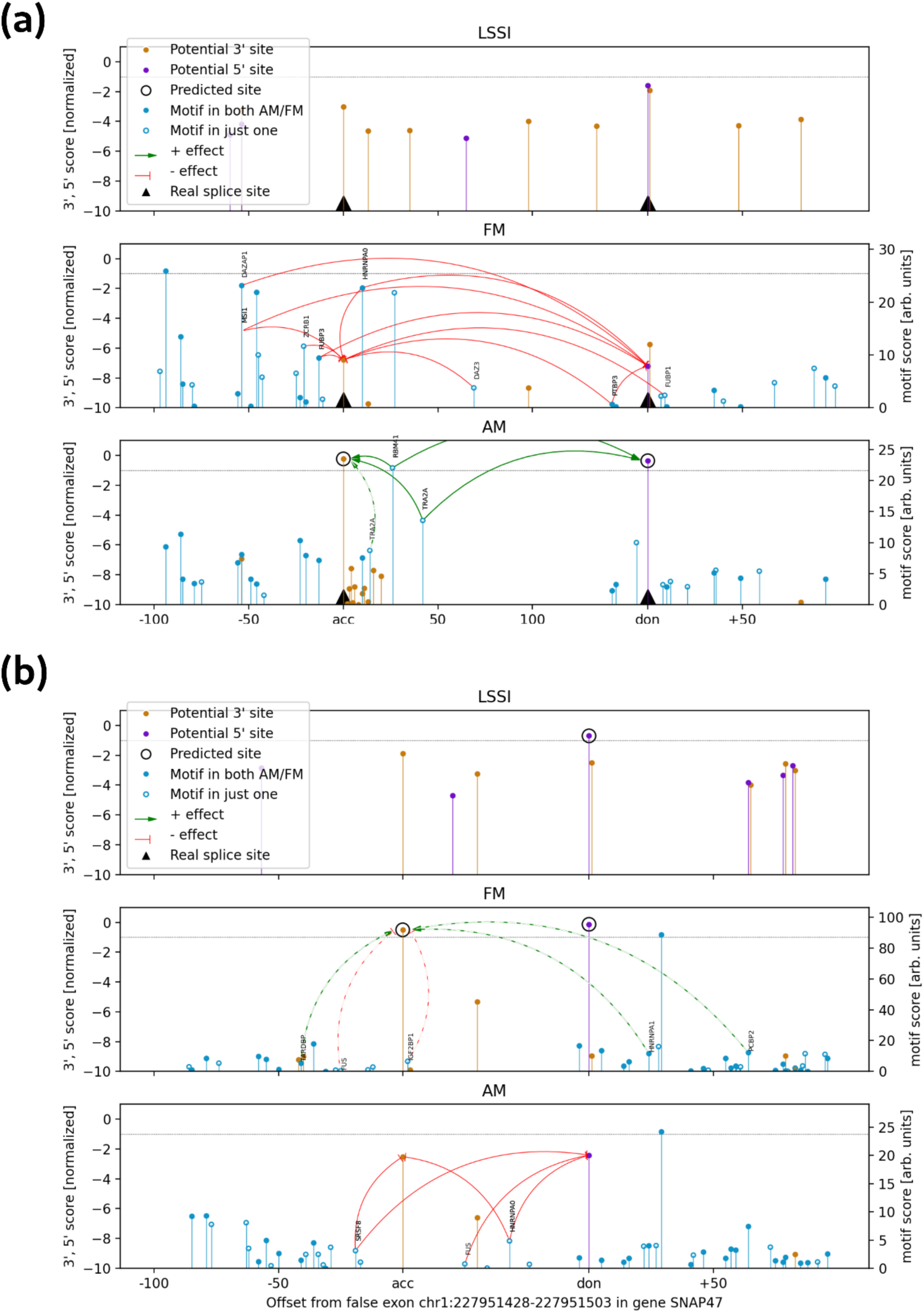
Examples of regulatory logic of FM and AM models for annotated and decoy exons. Predicted 3’SS and 5’SS are shown as orange and purple lollipops, respectively; height represents score, normalized so that –1 is the top-k threshold for the model for each class; only splice sites scored above –5 in the normalized score by at least one model are shown. Black triangles on the x-axis correspond to annotated splice sites, while circled predictions correspond to the predicted splice sites for each model. Motif predictions are shown as blue lollipops on the FM and AM plots, in relative affinity units. Filled circles represent motifs that are present in both models; empty circles are motifs that appear in only one. Solid arrows connect any motif to the corresponding splice site if an in-silico knockout of the motif increases the splice site prediction probability by 1.5-fold or more (green) or decreases it by a factor of ⅔ or less (red arrows). At least the top 5 effects are shown for each exon, using dashed arrows to indicate those that do not meet the cutoff for solid line.

We also analyzed the aggregate effects of individual motifs on splicing (Figure 7). For purposes of comparison, we show the motif effects predicted by our model for all hnRNP, SR and SR-related proteins studied. These protein classes were chosen as they have well-understood effects on constitutive splicing and cassette exons. In general, we observed a pattern in which hnRNP proteins typically have a negative effect in exons, a positive effect in introns, or both activities, with the opposite being true for SR and SR-related proteins. Such patterns were observed for 17 of the 23 RBP analyzed, with similar results obtained for FM models (Supplementary Figure 5), indicating that it is a property of our overall modular architecture. Exceptions to this pattern may result from noncanonical activities of some factors, e.g., SRSF10 is an atypical SR protein that can repress splicing in exons^37^, and this activity is identified by our model.

**Figure 7.**
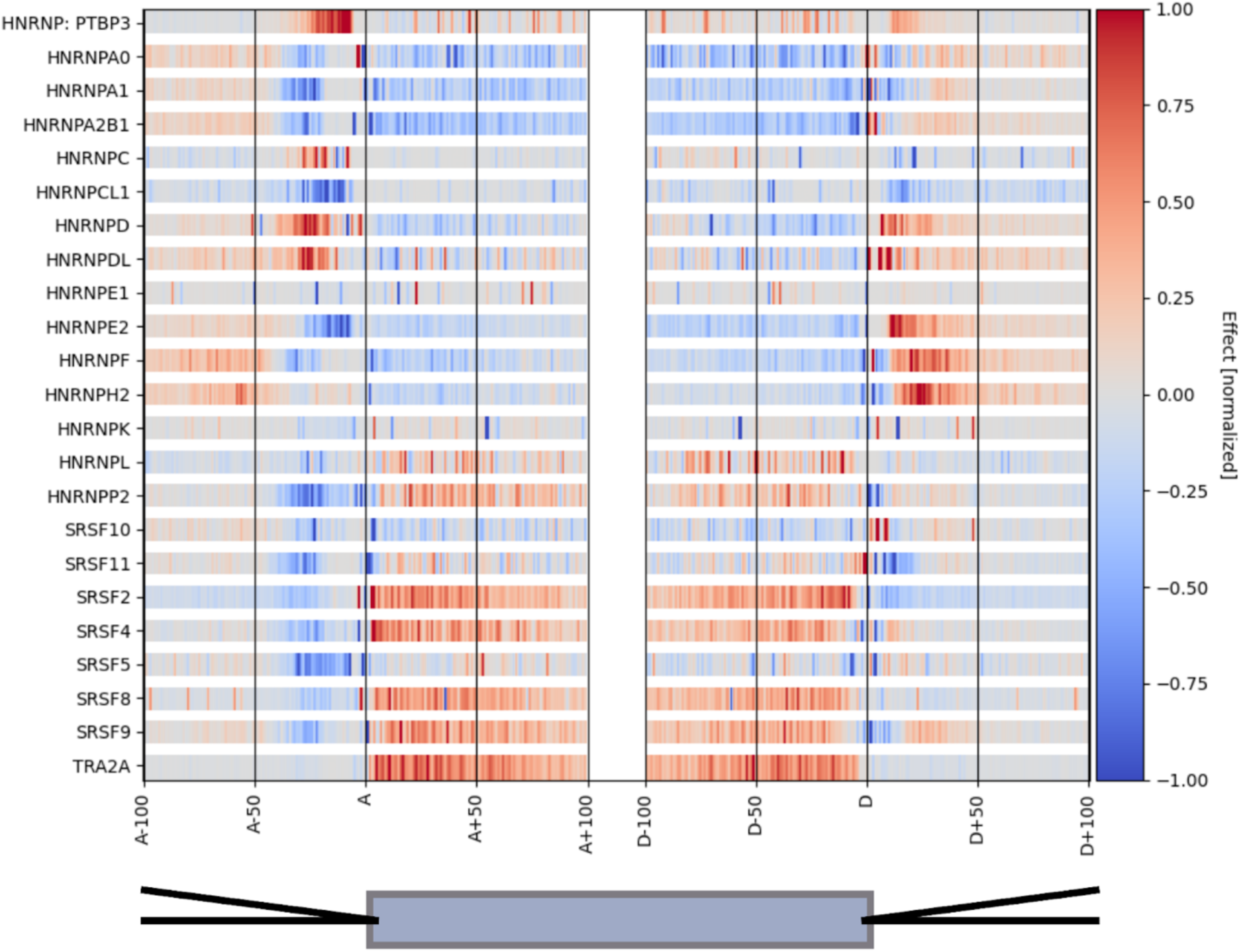
Inferred RNA maps for selected RBPs. To create this plot, we performed in-silico knockdowns on the binarized version of our model and measured changes to splicing. We then aggregated across several positions to create overall effects at each position in the meta-exon. The results here are normalized for each motif individually, with an absolute value of 1 corresponding to the 99th percentile across positions. This normalization enables us to see the RNA map for each motif. The x-axis here is in two parts, with the left half being relative to 3’ sites (A), and the right half being relative to 5’ sites (D); a schematic exon is shown at bottom.

A direct side-by-side comparison with the ENCODE RNA maps based on eCLIP and knockdown data^34^ yielded fairly strong overall concordance (Supplementary Figure 6). The small number of discrepant cases might result from technical biases in eCLIP or RBNS, or from ‘aliasing’ issues, where a motif associated with the RBP that was CLIPped and knocked down is also bound by another protein with different splicing regulatory activity. The inferred pattern of hnRNP and SR activity is consistent with many previous studies^1,38,39^. Our observations that 17 (18 counting SRSF10) of the 23 proteins exhibit the expected pattern of activity in cells subject to RBP knockdown supports that the Aggregator has learned splicing regulatory activities of many RBPs.

### Prediction of Other Classes of Splice Sites

The transcript annotations used in the majority of our model training and testing were taken from the SpliceAI ‘canonical’ gene annotations, which include a single annotation for each gene based on the principal transcript from that gene and therefore include predominantly constitutive exons. But most human genes contain additional alternative exons, and some contain evolutionarily new exons, both of which are likely to be more challenging to predict. Previously, it was reported that SpliceAI yields somewhat weaker predictions on alternative exons relative to constitutive exons^18^, and we observed a systematic shift toward lower accuracy for all models on alternative exons. While further study is needed, likely contributors to this pattern are that alternative exons have weaker splice sites and a regulatory element composition that is less biased toward positive-acting elements than constitutive exons (Figure 8)^40^.

**Figure 8.**
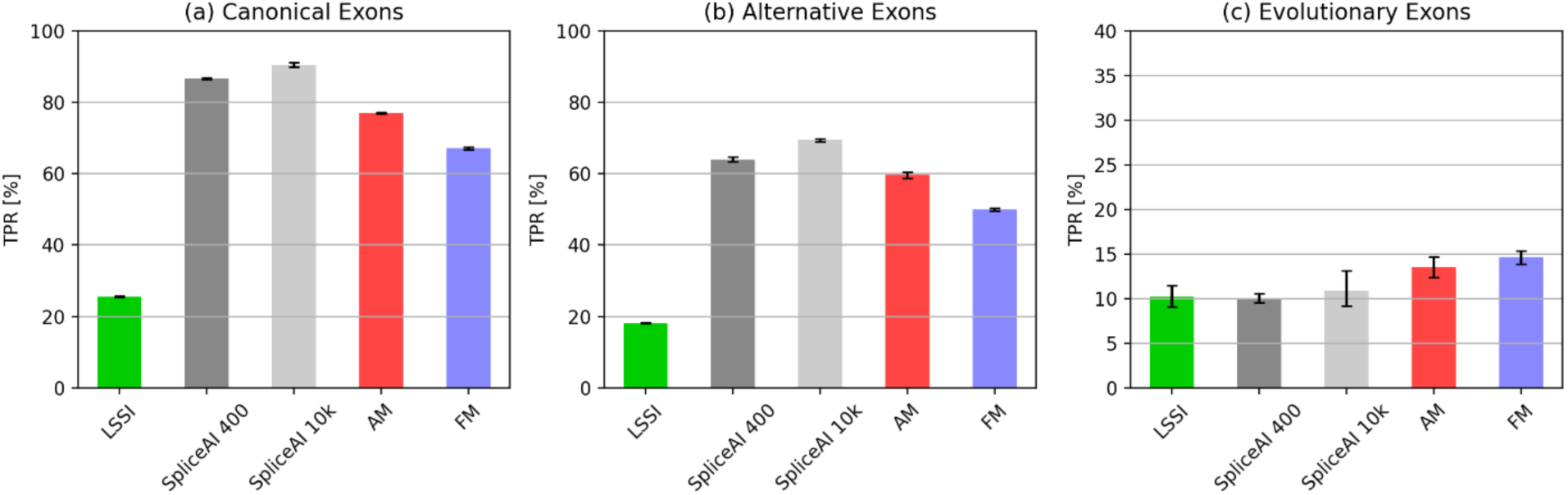
Accuracy for different classes of exons. Top-k accuracy of splice site prediction for models trained on canonical splice sites applied to (a) canonical exons; (b) alternatively spliced exons; and (c) evolutionarily new exons (note that the y-axis range was reduced for this panel to facilitate comparison). Here, the value of “k” used for top-k accuracy is the union of the numbers of splice sites across all 3 classes, rather than per splice site type: this approach ensures a consistent threshold across the three conditions. Error bars are 95% bootstrap intervals of the mean across 5 replicates.

We also analyzed a previously described set of evolutionarily new internal human exons (i.e. genomic segments spliced as exons in humans but not spliced as exons in rodents or other mammals), which we anticipated would be difficult to predict due to their weaker splice sites, frequent alternative splicing, and sequence features that are reminiscent of their intronic origins^41^. (The splice sites of alternative exons and evolutionarily new exons overlapped to only a small extent with the canonical splice site dataset (Supplementary Figure 7).) As expected, the splice sites surrounding evolutionarily novel exons proved far more challenging to predict for all models (Figure 8c). On this challenging task, our interpretable models (AM, FM) performed moderately better than others.

## Conclusions

Here we have shown that a neural model using core types of RNA elements known to contribute to splicing mechanisms can achieve fairly high accuracy and can learn binding and regulatory activities of RBPs. Indeed, we find that the AM models actually provide improved descriptions of the in vivo binding and regulatory activity of RBPs involved in splicing. Furthermore, our approach can infer RNA maps for splicing factors purely from in vitro binding data and the genome sequence, i.e. without requiring the CLIP and knockdown/RNA-seq data typically required for RNA map inference. We can also generate a putative regulatory landscape for any exon, which could have applications to genetic variant interpretation or design of therapeutic antisense oligonucleotides to perturb splicing. Furthermore, the general AM approach could potentially be applied outside of splicing to study other RNA- or DNA-binding proteins for which a defined regulatory activity and suitable training set are available.

## Availability of Code

The computer code needed to reproduce the analyses shown in this manuscript is available on GitHub at https://github.com/kavigupta/sam/tree/main/spliceai/Canonical

## Acknowledgments

We thank members of the Burge and Solar-Lezama groups for helpful discussions and Ben Blencowe and Manolis Kellis for helpful comments on the manuscript. This work was supported by grants from the NSF (to A. S-L.) and the NIH (to C. B. B.) and by a DOE-CSGF Fellowship (to K. M.).

## METHODS

### LSSI Architecture

We structure our LSSI models as simple feedforward neural networks that first map their input to 100 hidden neurons, then have 5 hidden 100-wide layers with ReLU activations, and then finally mapping to a prediction output. These models are trained on the same training dataset as our other models, using 10 epochs, a batch size of 150, and a learning rate of 10^-3^.

### Derivation of entropy bound

Here, we provide the derivation of the bound *H/L* ≤ *M[H(B(δ)) + δ η]*, where *B(x)* represents the Bernoulli distribution with probability *x*, *δ_i_* represents the sparsity of motif *i*, *η_i_* represents the entropy per nonzero value of channel *i*, and *η* represents the entropy in the overall distribution of activations across channels:

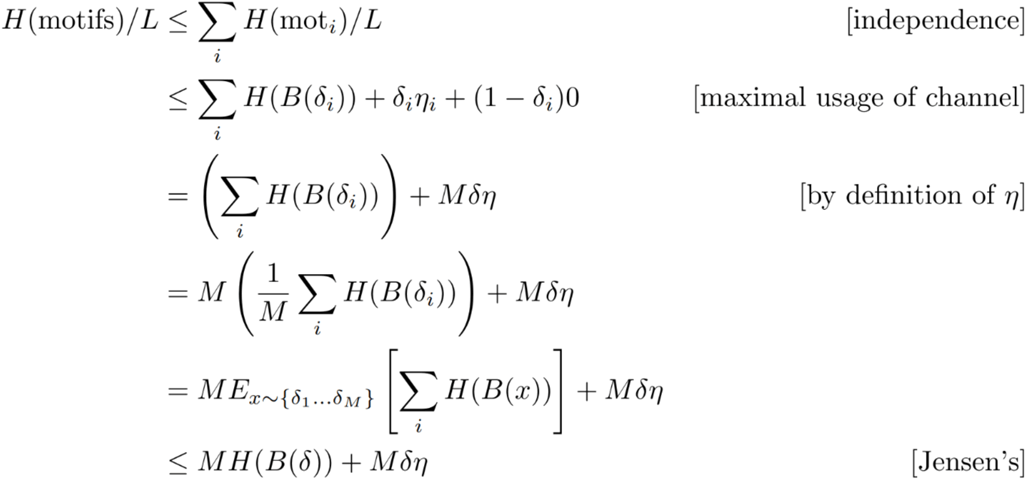

### Adjusted Motif Model

We compute the binding sites in the adjusted motif (AM) model by adjusting the scores for already relatively high-scoring FM binding sites, allowing *k* times as many FM sites as we will eventually use, allowing our AM model to filter out a subset of binding sites among plausible sites provided by the FM model, as follows:

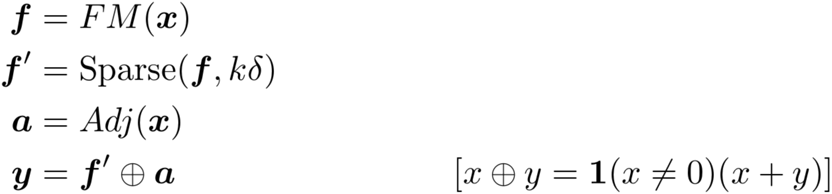

### Neural models trained on RBNS data

Given that the FMs are derived via the RBPamp algorithm from the RBNS dataset, we also tested the hypothesis that a neural version of motifs trained from the RBNS dataset could better represent protein binding. These neural motifs are also used as baselines to compare the performance on RBNS binding from AMs. We formulated the training task of the RBNS dataset as a binary classification task. Given a sequence from the RBNS dataset, we measure the accuracy of classification of the sequence being from the protein-bound or input RNA pool in the original RBNS experiment. Specifically, we select the sequence binding dataset for each protein with the highest 5-mer enrichment (“R value”) from the RBNS dataset. We trained our “Adjusted Motif” architecture on this dataset, but did not enforce sparsity. This becomes effectively just the sum of a PSAM signal and a learned neural model. This approach was used because it allows the learned model to have the advantage the AMs have of being able to take advantage of the PSAMs, but we do not want to constrain the model to be similar to PSAMs since it is being trained on the RBNS data directly.

We also separately trained 4-layer convolutional neural networks with two sliding window width choices, width=11 (NM11) and width=14 (NM14), in order to ensure that we are not too heavily tied to a single model architecture. Testing these models with a sparsity of 0.18% calibrated on the genome, we find that the NM14 and AM models improve on the RBNS dataset: gaining an improvement of 0.33% and 1.76% respectively, while the NM11 does not (it yielded a –0.02% change in performance). Our NFM models also improved on eCLIP, though the results were highly inconsistent. On average, they achieved about 6% of the improvement of the AM-E models, but their overall range of improvements across 4 replicates overlaps that of the AM.

### Analysis of MPRA data

In the case of the 5’SS MPRA data, the leftmost 3 bases in the targeted region do not have 10 bases of context to their left (as there are only 7 bases in the SD1 region). Since our AM models are 21-wide, they require 10 bases on either side of any site where we would predict a motif, we removed the leftmost 3 positions from consideration. For consistency, we did this for both the AMs and the FMs (it would not be strictly necessary for FMs, since they are only 11-wide).

### Analysis of exon classes

We consider both alternative and evolutionarily new exons whose splice sites were both located inside their respective gene boundaries as determined by the canonical dataset. If an exon’s associated gene was not included in the canonical dataset it was also excluded from consideration. The majority of the resulting splice sites were not included in the canonical dataset, though we found 76 splice sites in common between the canonical SS and the evolutionarily new SS, and around 180 SS in common between the alternative SS and the evolutionarily new SS, as shown in Fig. 7.

## Declaration of Interests

The authors declare no competing interests.

## Supplementary Text and Figures

### Entropy of nonzero motif values

To bound the number of bits of entropy in the nonzero motif binding sites, we run a binning experiment where we bin the nonzero motif binding sites into a certain number of bins based on rounding to the nearest k, then use the mean of each bin as the value for all elements of the bin. We then calculate both the empirical entropy of this distribution as well as the drop in accuracy when this rounding scheme is employed (Supplementary Figure 1). In general, we find that FMs have no entropy, whereas AMs have similar empirical entropies, both about 1.5-2b/activation. Both are below the 2b threshold that is needed for our 0.18% sparsity to translate into a sub-1.91b/nt entropy.

**Supplementary Figure 1.**
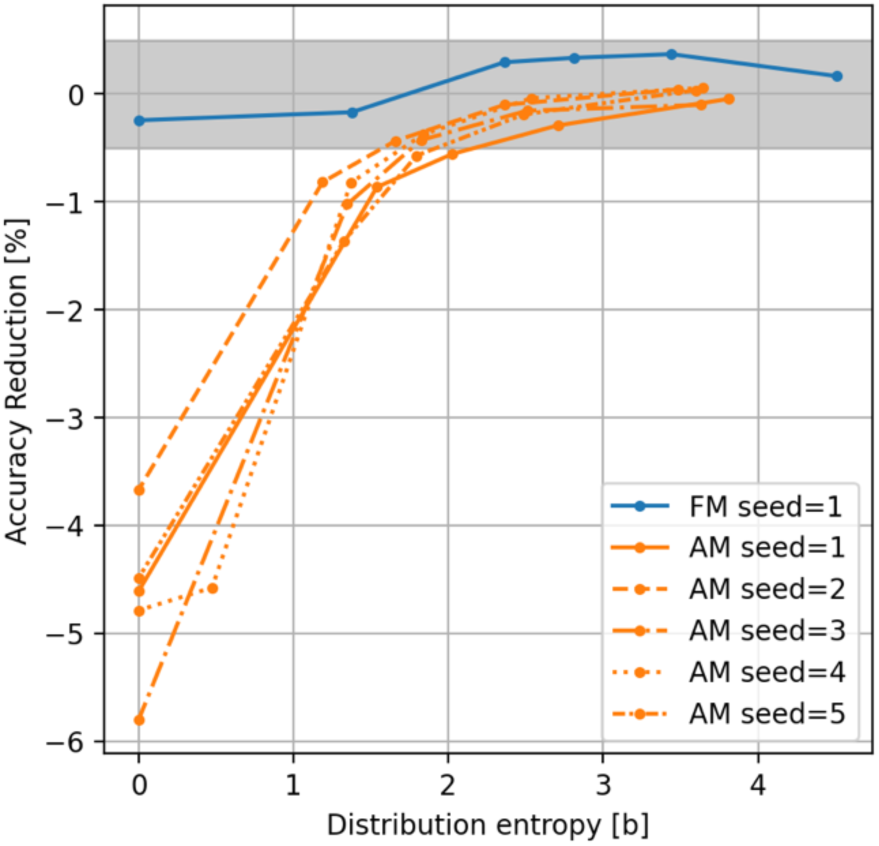
The effects of score binning on FM and AM model predictions. On the x-axis is the number of bits of entropy in the distribution, on the y axis is the reduction in accuracy from the original. We consider a reduction or increase of 0.5% to be within the noise region, as modifying a model in an open-loop format in any way will lead to some degradation in performance.

### Effect of Aggregator Architecture

While the focus of our paper is on the Motif model and sparsity bottleneck rather than the Aggregator, we feel that our Aggregator architecture (referred to here as attention + recurrence, or A+R) has some benefits in terms of encouraging the motif models to behave more like RBP binding predictors. We use the SpliceAI-10k model as an aggregator as a baseline in order to demonstrate these properties. The SpliceAI model is a very flexible convolutional model, and also has a slightly longer context window which allows it to perform somewhat better than our Aggregator on end-to-end splicing. However, our primary focus was on motif quality. To explore this question, we performed a module substitution experiment (Supplementary Figure 2). In this experiment, we extracted motif models from AMs trained with either the A+R or the SpliceAI model as aggregator, and then we paired these with either the A+R or SpliceAI aggregator trained on FMs. What we find is that regardless of the ultimate aggregator used, the Motif Models trained with the A+R aggregator outperform those trained with the SpliceAI aggregator. This observation indicates that our A+R aggregator improves overall motif quality and thus interpretability, though it’s end-to-end accuracy does not match that of SpliceAI.

**Supplementary Figure 2.**
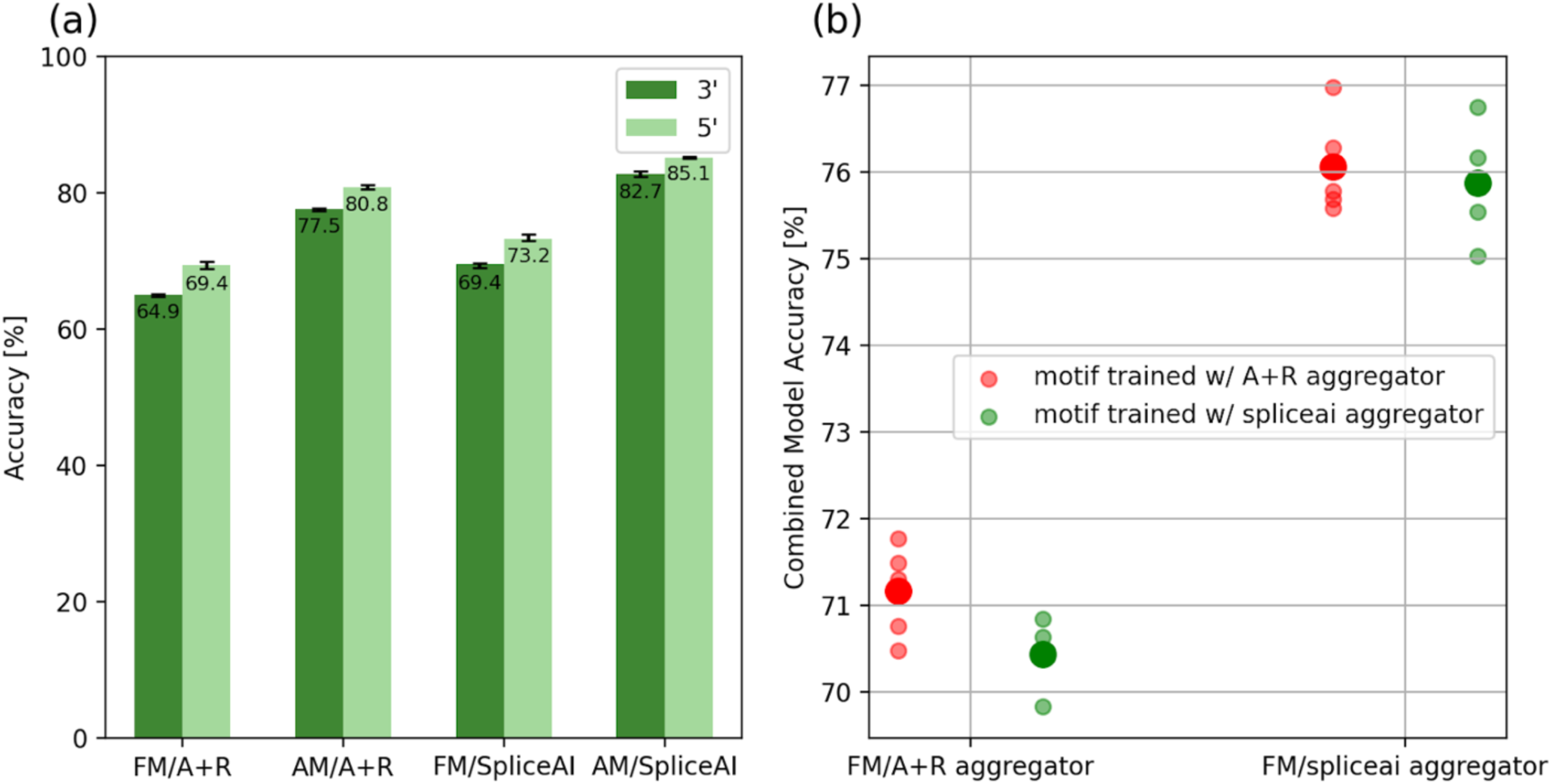
Effects of substitution of Aggregator architectures on performance. (a) End-to-end splicing results for FMs and AMs using both the A+R and SpliceAI aggregator. (b) Module Substitution Experiment results for combining A+R and SpliceAI-trained AM motif models with FM aggregators of both varieties.

### Motif Width

Our Adjusted Motif Models take in a context length of 21 nt. We find that this length – somewhat longer than typically used to model RBP binding – yields a pronounced improvement in performance. In this section, we contrast 21 nt models with 13 nt ones, length that more closely approximates the 11 nt used in the RBPamp PSAMs^27^. We note that shrinking our motifs from 21 to 13 reduces accuracy by ~1.7%. Below, we analyze several possibilities for what might case this difference.

*Motif architecture*. First, we examined the motif models themselves. The shorter AM is composed of a smaller number of Residual Units, and as such has a theoretically smaller computational capacity. However as seen in part (a) of Figure 3a, adding in additional layers that do not extend the width of this model does not improve performance, and in fact slightly reduces it.

*Secondary structure*. Another hypothesis is that RNA secondary structure can be more easily captured by longer motifs. To test this hypothesis, we provided the mean basepairing probability to the algorithm from a standard thermodynamic RNA secondary structure prediction algorithm. To do so we began with sequences of length 40 nt, adding 30 nt of context to either side, then used RNAFold^42^ on the 100 nt sequences, computing the probabilities of all base pairs in secondary structure. We then sum across to find the probability of any given base being bound in secondary structure, and discard the 30 nt on each side, leaving the more reliable center 40 nt. We find that this improves performance by about 0.4%, but this could provide some information about the sequence so is difficult to interpret. To properly control for this, we also provide an alternate source of information about the sequence, where we run the exact same procedure, but first swap all A and C bases in the sequence. This produces a similar kind of signal from an informational perspective, but is not in any way related to secondary structure. We find that there is no difference in the performance of this model and the model using the original secondary structure. This experiment suggests that capturing the overall secondary structure potential of a region is not particularly helpful in prediction and is therefore not likely to explain the advantage of wider motifs, though it doesn’t rule out that some other aspect of RNA secondary structure might be involved.

**Supplementary Figure 3.**
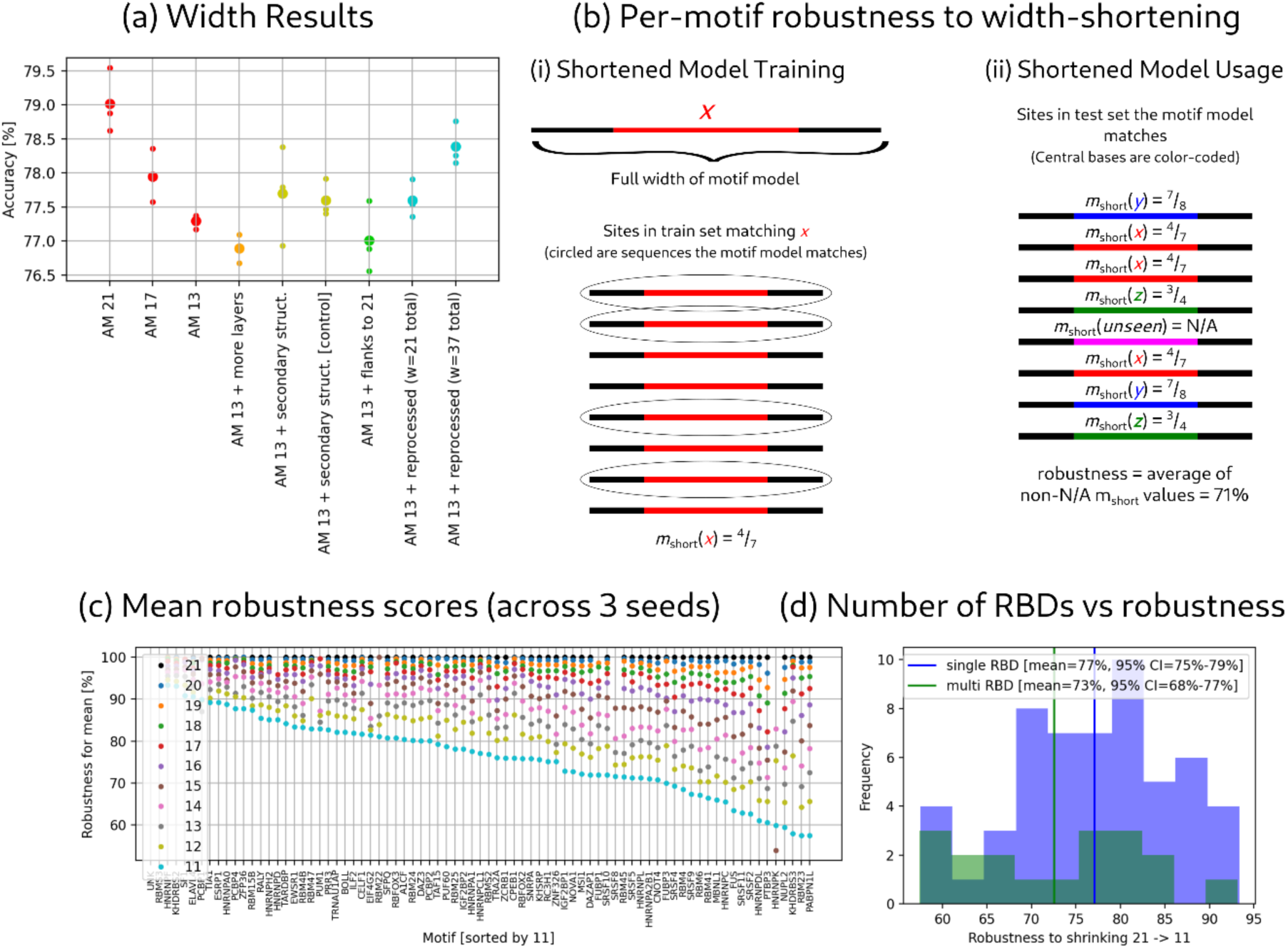
Motifs of width 21 are optimal overall. (a) Accuracy for several different motif models. In general, we find that greater width leads to greater accuracy, and this cannot be made up with more layers, secondary structure, a flank-included motif model, or a reprocessor layer. (b) (i) Shortened motif model training: for all core sequences of length w – a – b, we scan the training dataset (from the genome) and find all instances of matches, then find how many of them are at the center of matches of the full motif model. Our shortened model outputs this fraction. (ii) Testing: we then average the values of the shortened motif model on all matches of the motif model. (c) Motif width-reduction robustness values for a variety of widths, from 21 down to 11. Below 11 the numbers tend to drop far more quickly as we start clipping the FMs. The motifs here are ordered by their robustness at 11, and the values here are taken as a mean over several other motifs. (d) Histogram of single and multi-RBD motifs’ robustness to shrinkage. There is no apparent relationship between robustness to shrinkage and number of RBDs.

*Flanking sequence context*. We also considered the possibility that there is some broad flanking context that is useful in predicting whether an RBP will bind a site. For example, perhaps an RBP binds more frequently in AT-rich sequences over GC-rich sequences^26^. To explore this possibility, we trained a model that uses the core 13 nt AM, but adds in a 21-wide “flanking model”, which is effectively a PSAM but with the restriction that the ratios between the log affinities of each base are fixed at each position – in effect, requiring that the same preference be applied to each position, though with potentially different magnitudes. This approach also failed to increase accuracy, suggesting that increased width being more accurate is not closely related to general flanking base composition.

*Cooperative/competitive motifs*. Finally, we investigate the possibility that our longer motif models are picking up on competition between motifs – effectively, that the “21 wide motifs” actually represent a smaller motif that has learned to recognize other motifs in the same window. We investigate this using a Reprocessed Motif model, which takes the outputs of the 13-wide AM model and runs them through another convolutional network. When we make this network 9 wide, in order to make our overall width 21, we end up with slightly better performance (+0.3%), but less than that achieved by a 17-wide motif. Even when we make the Reprocessor layer 25 wide (total width of 37), we are unable to improve to the same performance as a 21-wide network. This result argues against the idea that other motifs are being picked up in the 21-wide window, but has the caveat that motifs outside our considered set might be learned.

### Robustness Scores

We also investigate the width per-motif by using our robustness to width-shortening method. We compute this for any number of bases removed from the left and the right by “training” a shortened motif model as depicted in Supplementary Figure 3b. In doing so, we compute for every shortened sequence the probability that it is part of a true motif binding site. We then “evaluate” this shortened motif model by computing the mean score on all true binding sites of the motif model on the test set. We then compute the true robustness metric at a certain length by taking the maximum over all available shortened windows. We plot a graph of this metric, averaged across three separate AM models, for all motifs and various widths. Using this metric, we can test another hypothesis, that our adjustment model is using a larger width in order to pick up on multi-RBD motifs. However, if this were the case, we would expect much lower robustness to shrinking among models with multiple RBDs, which was not observed (Supplementary Figure 3d).

Taken together, these experiments failed to yield a compelling explanation for the performance difference between 21 and 13 wide models. It is possible RBP motifs have longer binding sequences than previously thought, or some other factor we have not considered could be underlying this phenomenon.

#### eCLIP metric

To compute intron/exon-controlled eCLIP enrichment for a set of predicted motif binding sites, we performed the following procedure. First, we created a “control peaks set” that consisted of the same number of peaks per motif, but distributed at random throughout the test set. We then split up both the real and control peak sets into intronic and exonic sets based on whether they were entirely in the intron or exon (peaks that crossed a splice boundary were discarded). Next, we computed the fraction of eCLIPs under each of the 2 x 2 x 18 conditions (real/control, intron/exon, motif) that were covered by the given motif binding sites, where coverage was defined as containing a nonzero value. We then averaged across motifs to leave us with 2 x 2 values, the coverage_real,x_ and coverage_control,x_ for x in [intron, exon]. The final enrichment values produced are enirchment_x_ = coverage_real,x_ / coverage_control,x_.

The need for splitting up introns and exons comes from the well-known bias of eCLIP towards exonic peaks [cite], which might otherwise bias our results if, for example, the AMs were promoting exonic binding sites over intronic ones. Averaging across motifs was done to reduce the variance of the experiment. Finally, the control eCLIP set was done to ensure that we had an interpretable final result, enrichment, rather than coverage. This, however, did not make our metric insensitive to the sparsity of the motif binding sites; as less sparse motif binding sites would often get lower enrichment values as there was less room for them to provide signal.

As a result, we made sure to match the sparsity of the FMs exactly on each motif before taking the relative enrichment under intron and exon, which was computed as relative_enrichment_x_ = (enrichment_tested model,x_ – enrichment_FM,x_)/enrichment_FM,x_. Attempting to somehow take a weighted average of these conditions was problematic as introns are far larger but exons are generally closer to splice sites and thus each nucleotide is on average more important in splicing. Thus, we opted instead to present them separately.

#### Non-interpretability of SpliceAI model

Initially, we attempted to extract a directly interpretable model from SpliceAI. However, we found that considering sequence perturbations was not clarifying, as it appears that the model uses information widely distributed across the sequence. To demonstrate this, we performed the following experiment, using SpliceAI-400 for ease of calculation. In the experiment, we considered strong true positive splice sites: those which SpliceAI predicts correctly with at least 10% margin over the top-k threshold. For each given splice site, we computed a “perturbation swing” at every input base, defined as the maximum absolute difference in the predicted probability of the splice site that can result from substituting the given base, testing each of the three alternate bases. We then varied a threshold on the perturbation swing and randomized all bases whose swing values were below this threshold, in essence randomizing the portions of the sequence that are considered less important by SpliceAI.

The results of this experiment, averaged across several different splice sites, are depicted in Supplementary Figure 4a. At a swing threshold of even as low as 1%, we find that 60% of the sequence falls below this threshold, and that randomizing these bases leads to a 25% decrease in accuracy. Since we selected for robust splice sites with probability at least 10% above the threshold, this indicates that many positions are being used in tandem to produce the overall signal for splicing. Thus, single perturbations are insufficient to explain SpliceAI predictions in general, even for SpliceAI-400.

Additionally, we considered whether single base perturbations are sufficient to infer motif locations, potentially obviating the need for an explicit motif model. To test this idea, we determined the distance from the nearest motif (using the FM model with density 0.18% to define the motif positions; this is effectively the RBNS motifs) and in each distance category we computed the mean perturbation swing. We excluded the 40 bases closest to the splice site of interest to ensure that the core splice motif is not contaminating the results. Surprisingly, we found that distance from the nearest motif center positively rather than negatively correlated with perturbation swing, as seen in Supplementary Figure 4(b)(i). This was true even controlling for LSSI sites (splice site motifs scored with log probability −10 or higher by the LSSI model), by looking only at positions that are at least 20 nt away from the nearest LSSI site. We can conclude that while SpliceAI presumably uses information about SREs to make splicing predictions, it clearly is using other information to a greater degree, leading to its predictions being uninterpretable from the point of view of the SREs considered here, which include motifs for many canonical splicing factors of the hnRNP, SR and SR-related protein classes likely representing a substantial portion of splicing regulatory motifs. As a check of our metric, we show in Supplementary Figure 4 (b)(ii-iii) that this property is unique to SpliceAI and does not apply to either of our models, both of which show the opposite trend, starting with similar absolute swing magnitudes, with perturbation swing declining rather than increasing as we move further from the nearest motif.

**Supplementary Figure 4.**
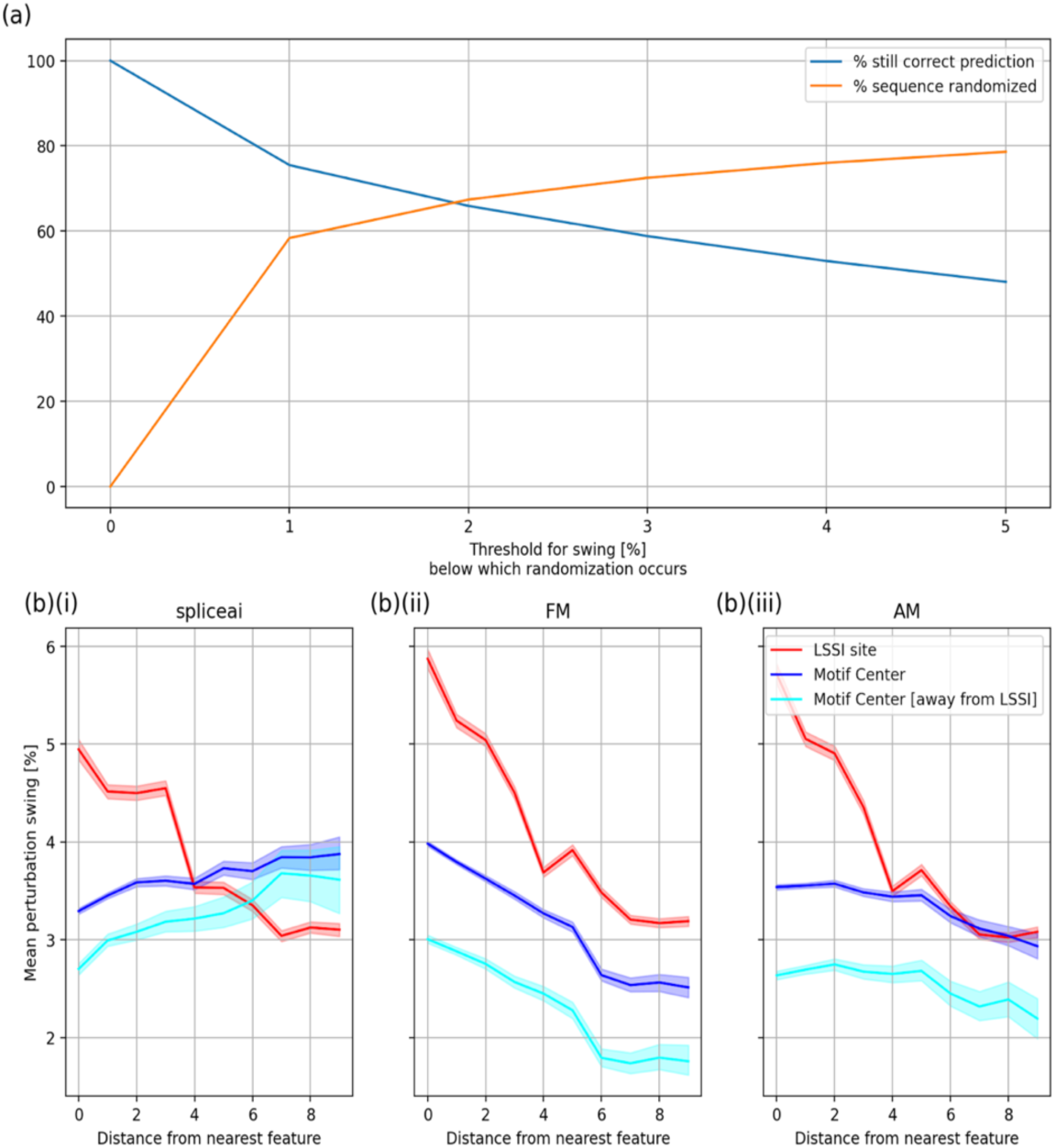
Non-interpretability of SpliceAI. (a) Accuracy of SpliceAI-400 applied to partially randomized sequences, as described in text. (b) Mean perturbation in score resulting from in silico mutations at different distances from LSSI sites (splice site motifs, red), RBP motif centers (dark blue), or RBP motif centers located at least 40 nt from LSSI sites (light blue) The distance from the nearest feature is shown on the x-axis (in nt), and the mean perturbation swing on the y-axis.

#### Use of RNACompete PSAMs

To ensure that our results are not specific to RBNS PSAMs, we also consider RNACompete and a hybrid model where we merge the RNACompete and RBNS sets, preferring RBNS motifs when we have both available. We sourced RNACompete PSAMs from Ray et al^43^, filtering for vertebrate RBPs. We consider the same RBP from different species to be different motifs. As such, we end up with a set of 97 motifs in the RNACompete set and 143 in the hybrid set. We calibrate our sparsity to have an approximately equivalent entropy as 79 motifs at 0.18% sparsity, and achieve an AM accuracy of 78.7% on RNACompete and 79.0% on our hybrid set. This demonstrates that our method is not specific to the RBNS motifs and works across different motif sets, but since these are larger motif sets and do not produce a change in performance, we continue to work with the smaller 79 motif RBNS set.

#### Additional examples of cases where AMs and FMs differ in prediction

Examples are provided at <u>this folder</u>. Several examples of situations where AMs predicted an exon correctly and FMs did not, on human, mouse, and fly. Examples for fly are scaled to (–2.5, 0) rather than (–10, 0) since the thresholds seem a bit lower for this organism. The accuracy gap between FM and AM is also lower for fly, leading to these selected exons being somewhat less representative than similar examples chosen for human or mouse. Exons are selected for size, LSSI incorrectness, FM incorrectness, AM correctness, the presence of TRA2A in the exon (for positive human examples; we did so because this motif has a well-understood and strong effect so it serves as a sanity check for the correctness of the splicing mechanism) and being on the positive strand (for convenience when viewing on a genome browser).

**Supplementary Figure 5.**
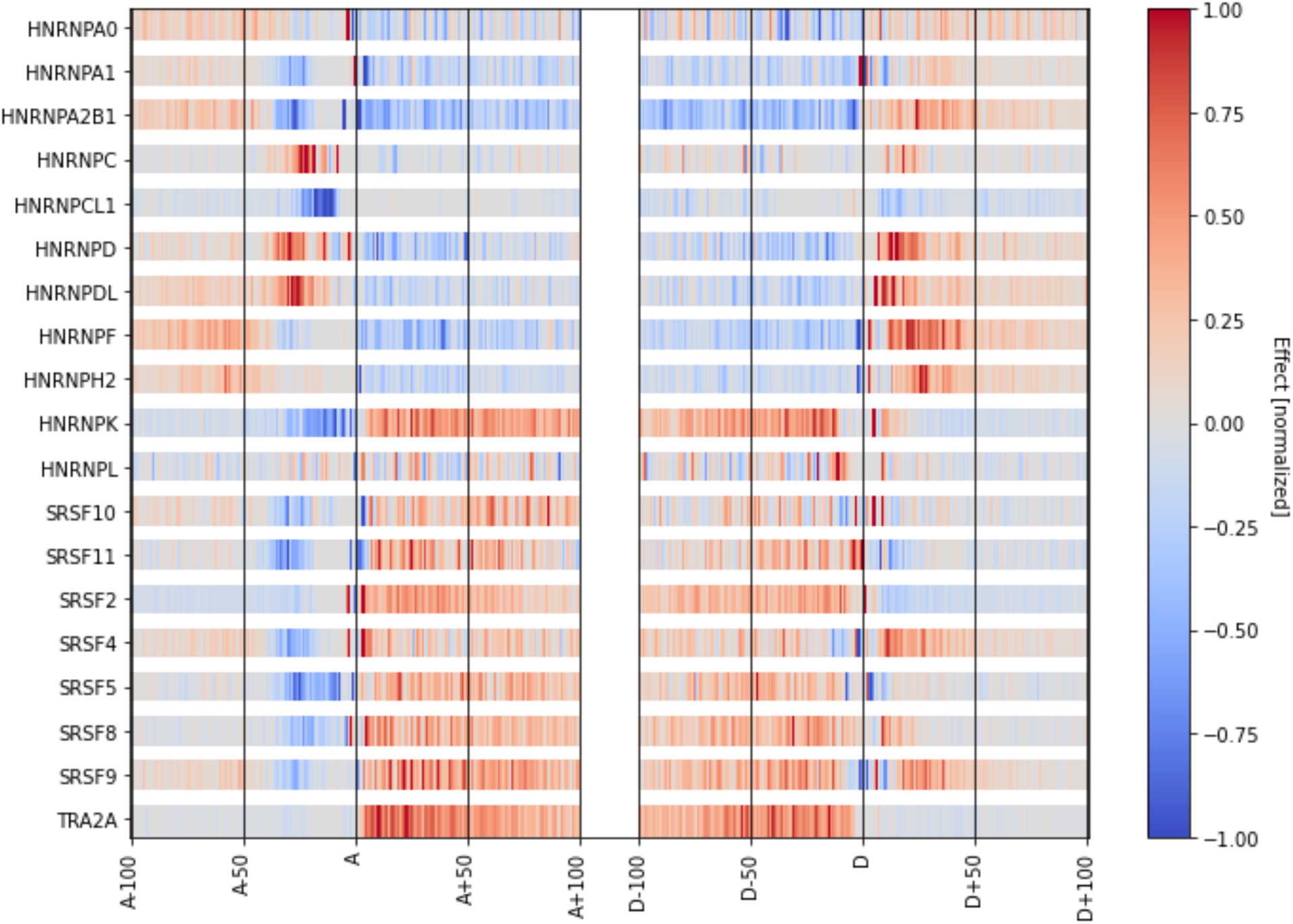
RNA maps using FM models. Similar to Figure 7. Data averaged over 5 FM replicates. Like AMs, the effect is broadly consistent with literature, with potential exceptions for HNRNPCL1, HNRNPK, and HNRNPL.

**Supplementary Figure 6.**
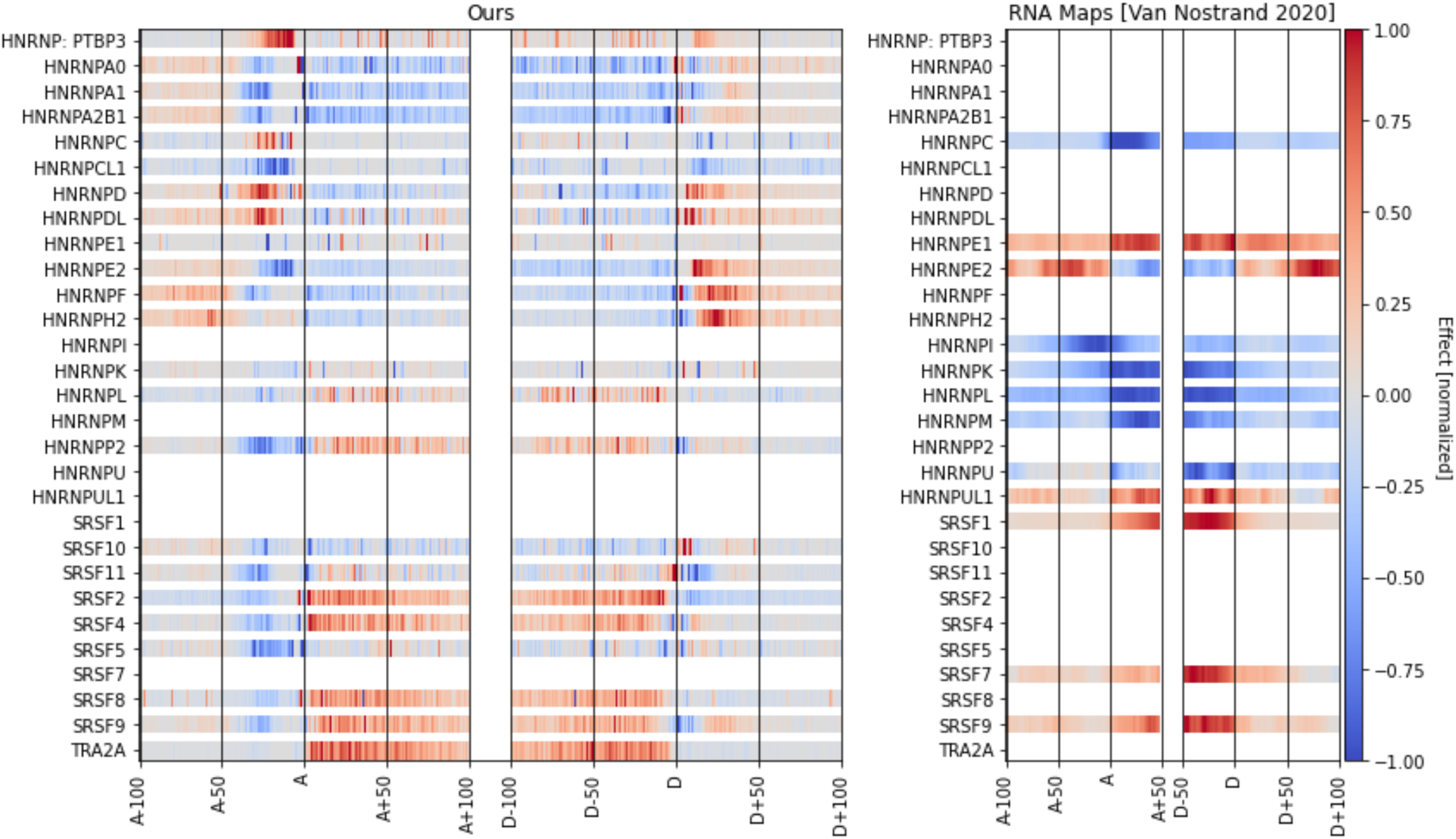
Comparison of SAM-based RNA maps to eCLIP/knockdown-based maps. In order to be consistent, we took only the parts of the maps that corresponded to the region near each splice site. Additionally, we combined the two maps by subtracting excluded - included. Finally, we performed the same normalization procedure we performed on our own motifs. In general, the Van Nostrand maps are smoother, with effects often bleeding across splice junction boundaries; this may result from their use of eCLIP peaks (which often have widths of 50-100+ nt) versus our pointwise representations of motifs.

**Supplementary Figure 7.**
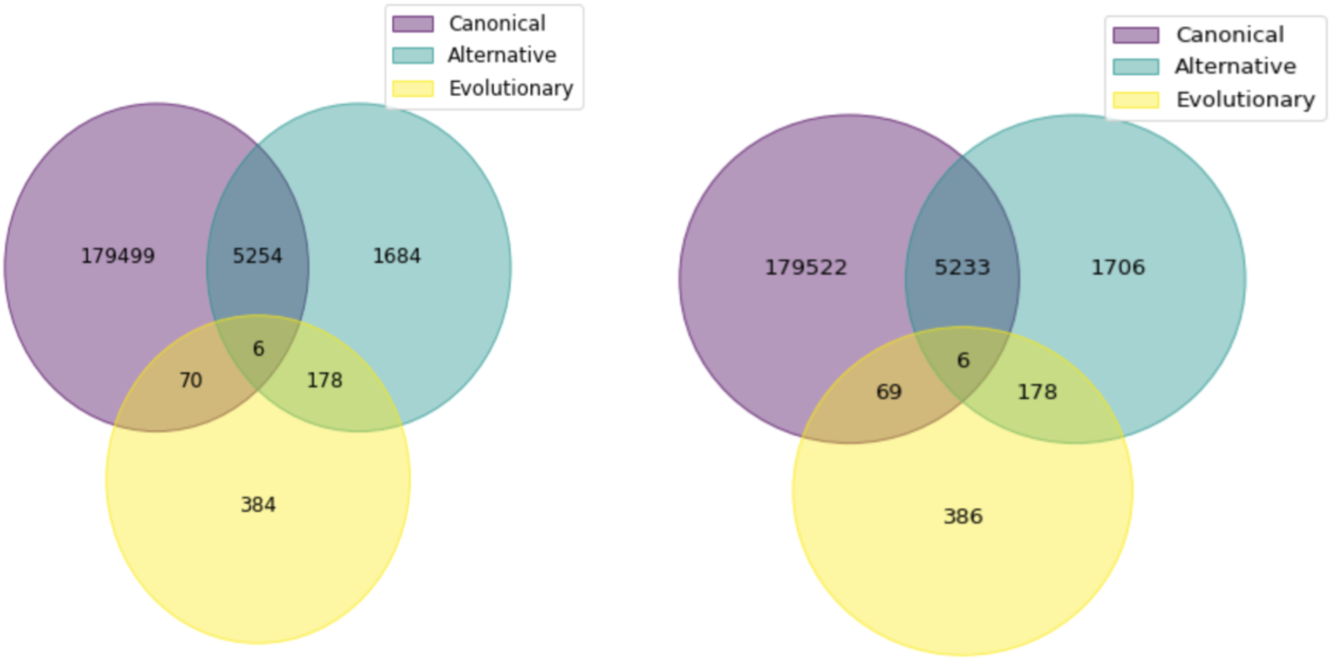
Overlaps between exon class datasets. The left and right figures represent the set distribution of 3’SS and 5’SS in each dataset, respectively.

## Notes

### Competing Interest Statement

The authors have declared no competing interest.

## REFERENCES

1. Lee, Y., and Rio, D.C. (2015). Mechanisms and Regulation of Alternative Pre-mRNA Splicing. Annu Rev Biochem 84, 291–323. 10.1146/annurev-biochem-060614-034316.

2. Burge, C., and Karlin, S. (1997). Prediction of complete gene structures in human genomic DNA. J Mol Biol 268, 78–94. 10.1006/jmbi.1997.0951.

3. Shapiro, M.B., and Senapathy, P. (1987). RNA splice junctions of different classes of eukaryotes: sequence statistics and functional implications in gene expression. Nucleic Acids Res 15, 7155–7174. 10.1093/nar/15.17.7155.

4. Pertea, M., Lin, X., and Salzberg, S.L. (2001). GeneSplicer: a new computational method for splice site prediction. Nucleic Acids Res 29, 1185–1190. 10.1093/nar/29.5.1185.

5. Yeo, G., and Burge, C.B. (2004). Maximum entropy modeling of short sequence motifs with applications to RNA splicing signals. J Comput Biol 11, 377–394. 10.1089/1066527041410418.

6. Lim, L.P., and Burge, C.B. (2001). A computational analysis of sequence features involved in recognition of short introns. Proceedings of the National Academy of Sciences of the United States of America 98, 11193–11198. 10.1073/pnas.201407298.

7. Wang, Z., Rolish, M.E., Yeo, G., Tung, V., Mawson, M., and Burge, C.B. (2004). Systematic identification and analysis of exonic splicing silencers. Cell 119, 831–845. 10.1016/j.cell.2004.11.010.

8. Barash, Y., Calarco, J.A., Gao, W., Pan, Q., Wang, X., Shai, O., Blencowe, B.J., and Frey, B.J. (2010). Deciphering the splicing code. Nature 465, 53–59. 10.1038/nature09000.

9. Xiong, H.Y., Barash, Y., and Frey, B.J. (2011). Bayesian prediction of tissue-regulated splicing using RNA sequence and cellular context. Bioinformatics 27, 2554–2562. 10.1093/bioinformatics/btr444.

10. Leung, M.K., Xiong, H.Y., Lee, L.J., and Frey, B.J. (2014). Deep learning of the tissue-regulated splicing code. Bioinformatics 30, i121–129. 10.1093/bioinformatics/btu277.

11. Jha, A., Gazzara, M.R., and Barash, Y. (2017). Integrative deep models for alternative splicing. Bioinformatics 33, i274–i282. 10.1093/bioinformatics/btx268.

12. Bretschneider, H., Gandhi, S., Deshwar, A.G., Zuberi, K., and Frey, B.J. (2018). COSSMO: predicting competitive alternative splice site selection using deep learning. Bioinformatics 34, i429–i437. 10.1093/bioinformatics/bty244.

13. Leman, R., Gaildrat, P., Le Gac, G., Ka, C., Fichou, Y., Audrezet, M.P., Caux-Moncoutier, V., Caputo, S.M., Boutry-Kryza, N., Leone, M., et al. (2020). Novel diagnostic tool for prediction of variant spliceogenicity derived from a set of 395 combined in silico/in vitro studies: an international collaborative effort. Nucleic Acids Res 48, 1600–1601. 10.1093/nar/gkz1212.

14. Mort, M., Sterne-Weiler, T., Li, B., Ball, E.V., Cooper, D.N., Radivojac, P., Sanford, J.R., and Mooney, S.D. (2014). MutPred Splice: machine learning-based prediction of exonic variants that disrupt splicing. Genome Biol 15, R19. 10.1186/gb-2014-15-1-r19.

15. Wainberg, M., Alipanahi, B., and Frey, B. (2016). Does conservation account for splicing patterns? BMC Genomics 17, 787. 10.1186/s12864-016-3121-4.

16. Cheng, J., Nguyen, T.Y.D., Cygan, K.J., Celik, M.H., Fairbrother, W.G., Avsec, Z., and Gagneur, J. (2019). MMSplice: modular modeling improves the predictions of genetic variant effects on splicing. Genome Biol 20, 48. 10.1186/s13059-019-1653-z.

17. Wang, R., Wang, Z., Wang, J., and Li, S. (2019). SpliceFinder: ab initio prediction of splice sites using convolutional neural network. BMC Bioinformatics 20, 652. 10.1186/s12859-019-3306-3.

18. Jaganathan, K., Kyriazopoulou Panagiotopoulou, S., McRae, J.F., Darbandi, S.F., Knowles, D., Li, Y.I., Kosmicki, J.A., Arbelaez, J., Cui, W., Schwartz, G.B., et al. (2019). Predicting Splicing from Primary Sequence with Deep Learning. Cell 176, 535–548 e524. 10.1016/j.cell.2018.12.015.

19. Cheng, J., Celik, M.H., Kundaje, A., and Gagneur, J. (2021). MTSplice predicts effects of genetic variants on tissue-specific splicing. Genome Biol 22, 94. 10.1186/s13059-021-02273-7.

20. Zeng, T., and Li, Y.I. (2022). Predicting RNA splicing from DNA sequence using Pangolin. Genome Biol 23, 103. 10.1186/s13059-022-02664-4.

21. Liao, S.E., Sudarshan, M., and Regev, O. (2022). Machine learning for discovery: deciphering RNA splicing logic. bioRxiv, 2022.2010. 2001.510472.

22. Chen, Z., Bei, Y., and Rudin, C. (2020). Concept whitening for interpretable image recognition. Nature Machine Intelligence 2, 772–782.

23. Zarlenga, M.E., Barbiero, P., Ciravegna, G., Marra, G., Giannini, F., Diligenti, M., Shams, Z., Precioso, F., Melacci, S., and Weller, A. Concept Embedding Models: Beyond the Accuracy-Explainability Trade-Off.

24. Aytar, Y., Vondrick, C., and Torralba, A. (2017). See, hear, and read: Deep aligned representations. arXiv preprint arXiv:1706.00932.

25. Desjardins, G., Courville, A., and Bengio, Y. (2012). Disentangling factors of variation via generative entangling. arXiv preprint arXiv:1210.5474.

26. Dominguez, D., Freese, P., Alexis, M.S., Su, A., Hochman, M., Palden, T., Bazile, C., Lambert, N.J., Van Nostrand, E.L., Pratt, G.A., et al. (2018). Sequence, Structure, and Context Preferences of Human RNA Binding Proteins. Mol Cell 70, 854–867 e859. 10.1016/j.molcel.2018.05.001.

27. Jens, M., McGurk, M., Bundschuh, R., and Burge, C. (2022). RBPamp: Quantitative Modeling of Protein-RNA Interactions in vitro Predicts in vivo Binding. bioRxiv, 2022.2011.2008.515616.

28. Ray, D., Kazan, H., Chan, E.T., Pena Castillo, L., Chaudhry, S., Talukder, S., Blencowe, B.J., Morris, Q., and Hughes, T.R. (2009). Rapid and systematic analysis of the RNA recognition specificities of RNA-binding proteins. Nat Biotechnol 27, 667–670. 10.1038/nbt.1550.

29. Tishby, N., Pereira, F.C., and Bialek, W. (2000). The information bottleneck method. arXiv preprint physics/0004057.

30. Gupta, K., Bastani, O., and Solar-Lezama, A. (2023). SPARLING: Learning Latent Representations with Extremely Sparse Activations. arXiv preprint arXiv:2302.01976.

31. Yang, Z., Yang, D., Dyer, C., He, X., Smola, A., and Hovy, E. (2016). Hierarchical attention networks for document classification. pp. 1480–1489.

32. Weidmann, C.A., Qiu, C., Arvola, R.M., Lou, T.F., Killingsworth, J., Campbell, Z.T., Tanaka Hall, T.M., and Goldstrohm, A.C. (2016). Drosophila Nanos acts as a molecular clamp that modulates the RNA-binding and repression activities of Pumilio. Elife 5. 10.7554/eLife.17096.

33. Malki, I., Liepina, I., Kogelnik, N., Watmuff, H., Robinson, S., Lightfoot, A., Gonchar, O., Bottrill, A., Fry, A.M., and Dominguez, C. (2022). Cdk1-mediated threonine phosphorylation of Sam68 modulates its RNA binding, alternative splicing activity and cellular functions. Nucleic Acids Res 50, 13045–13062. 10.1093/nar/gkac1181.

34. 34. Van Nostrand, E.L., Freese, P., Pratt, G.A., Wang, X., Wei, X., Xiao, R., Blue, S.M., Chen, J.Y., Cody, N.A.L., Dominguez, D., et al. (2020). A large-scale binding and functional map of human RNA-binding proteins. Nature 583, 711–719. 10.1038/s41586-020-2077-3.

35. Findlay, S.D., Romo, L., and Burge, C.B. (2022). Quantifying negative selection in human 3ʹ UTRs uncovers constrained targets of RNA-binding proteins. bioRxiv, 2022.2011.2030.518628.

36. Rosenberg, A.B., Patwardhan, R.P., Shendure, J., and Seelig, G. (2015). Learning the sequence determinants of alternative splicing from millions of random sequences. Cell 163, 698–711. 10.1016/j.cell.2015.09.054.

37. Feng, Y., Chen, M., and Manley, J.L. (2008). Phosphorylation switches the general splicing repressor SRp38 to a sequence-specific activator. Nat Struct Mol Biol 15, 1040–1048. 10.1038/nsmb.1485.

38. Erkelenz, S., Mueller, W.F., Evans, M.S., Busch, A., Schoneweis, K., Hertel, K.J., and Schaal, H. (2013). Position-dependent splicing activation and repression by SR and hnRNP proteins rely on common mechanisms. Rna 19, 96–102. 10.1261/rna.037044.112.

39. Shen, M., and Mattox, W. (2012). Activation and repression functions of an SR splicing regulator depend on exonic versus intronic-binding position. Nucleic Acids Res 40, 428–437. 10.1093/nar/gkr713.

40. Garg, K., and Green, P. (2007). Differing patterns of selection in alternative and constitutive splice sites. Genome Res 17, 1015–1022. 10.1101/gr.6347907.

41. Merkin, J.J., Chen, P., Alexis, M.S., Hautaniemi, S.K., and Burge, C.B. (2015). Origins and impacts of new mammalian exons. Cell Rep 10, 1992–2005. 10.1016/j.celrep.2015.02.058.

42. Lorenz, R., Bernhart, S.H., Honer Zu Siederdissen, C., Tafer, H., Flamm, C., Stadler, P.F., and Hofacker, I.L. (2011). ViennaRNA Package 2.0. Algorithms Mol Biol 6, 26. 10.1186/1748-7188-6-26.

43. Ray, D., Kazan, H., Cook, K.B., Weirauch, M.T., Najafabadi, H.S., Li, X., Gueroussov, S., Albu, M., Zheng, H., Yang, A., et al. (2013). A compendium of RNA-binding motifs for decoding gene regulation. Nature 499, 172–177. 10.1038/nature12311.

